# Cell size and selection for stress-induced cell fusion in unicellular eukaryotes

**DOI:** 10.1101/2024.08.19.608569

**Authors:** Xiaoyuan Liu, Jonathan W. Pitchford, George W. A. Constable

**Author notes:** (XL).

## Abstract

In unicellular organisms, sexual reproduction typically begins with the fusion of two cells (plasmogamy) followed by the fusion of their two haploid nuclei (karyogamy) and finally meiosis. Most work on the evolution of sexual reproduction focuses on the benefits of the genetic recombination that takes place during meiosis. However, the selection pressures that may have driven the early evolution of binary cell fusion, which sets the stage for the evolution of karyogamy by bringing nuclei together in the same cell, have seen less attention. In this paper we develop a model for the coevolution of cell size and binary cell fusion rate. The model assumes that larger cells experience a survival advantage from their larger cytoplasmic volume. We find that under favourable environmental conditions, populations can evolve to produce larger cells that undergo obligate binary cell fission. However, under challenging environmental conditions, populations can evolve to subsequently produce smaller cells under binary cell fission that nevertheless retain a survival advantage by fusing with other cells. The model thus parsimoniously recaptures the empirical observation that sexual reproduction is typically triggered by adverse environmental conditions in many unicellular eukaryotes and draws conceptual links to the literature on the evolution of multicellularity.

**Author summary:** Sexual reproduction is commonly observed, both in eukaryotic microorganisms and in higher multicellular organisms. Sex has evolved despite numerous apparent costs, including investment in finding a partner and the energetic requirements of sexual reproduction. Binary cell fusion is a process that sets the stage for sexual reproduction by bringing nuclei from different cells into contact. Here, we provide a mathematical explanation of the advantage conferred by binary cell fusion due to increased cell mass. We show that when unicellular organisms have the option to invest in either cell fusion or cell mass, they can evolve to fuse together as rapidly as possible in the face of adverse environments, instead of increasing their mass. These results are consistent with the empirical observation that sexual reproduction is often triggered by environmental stress in unicellular eukaryotes. Our results imply an advantage to cell fusion, which helps to shed light on the early evolution of sexual reproduction itself.

## Introduction

Although the details of the early evolution of sexual reproduction in the last common eukaryotic common ancestor (LECA) are shrouded in mystery, it is argued that the emergence of eukaryotic sex began with the evolution of cell–cell fusion and meiosis [1] in an archaeal ancestor [2, 3]. This step can be further broken down into the evolution of binary cell fusion, the one spindle apparatus, homologous pairing and chiasma, and finally reduction, division and syngamy [4]. The vast majority of theoretical studies investigating the evolution of sexual reproduction have focused on later stages of this evolutionary trajectory, namely the conditions that give rise to a selective pressure for genetic recombination [5–8]. However, comparatively few studies have investigated the selective pressures that may have first given rise to binary cell fusion, which may have facilitated the evolution of a host of other eukaryotic traits [9], including the homologous pairing and meiotic recombination, by bringing nuclei together in the same cell.

Hypotheses for the evolution of binary cell fusion often rely on hybrid fitness advantage. It has been suggested that selection for cell–cell fusions might have initially been driven by “selfish” transposons and plasmids [10–12], or negative epistatic interactions between mitochondrial mutations [13, 14]. However, once a heterokaryotic cell has been formed (binucleate with nuclei from both parental cells), the advantage of hybrid vigor and the masking of deleterious mutations could lead to the maintenance of cell fusion [4]. Such benefits are required to alleviate costs to cell-fusion, which include selfish extra-genomic elements in the cytoplasm [15] and cytoplasmic conflict [16, 17].

In these previous studies on the evolution of binary cell fusion, the effect of changing environmental conditions is not considered. However, in many extant unicellular organisms, binary cell fusion (and the karyogamy and genetic recombination that follow) occur in response to challenging environmental conditions [18] such as starvation (*Tetrahymena* [19, 20]) and depleted nitrogen levels (*Chlamydomonas reinhardtii* [21] and *Saccharomyces pombe* [22]). Meanwhile in benign conditions with abundant resources these species reproduce asexually using binary cell fission. The mechanisms that drive selection for genetic recombination under challenging environmental conditions are well-studied [23]; recombination can facilitate adaptation to a novel environment [24, 25] and evolving to engage in more sex when fitness is low (fitness-associated sex) can allow an organism to maximise the advantages of sex while minimising the costs [26, 27]. However, this focus on the benefits of recombination leaves space to ask whether binary cell fusion itself could be selected for as a stress response, even in the absence of any genetic advantages.

In this paper, we do not account for the genetic factors discussed above. Instead, we focus only on how the survival advantage associated with increasing cytoplasmic volume might select for binary cell fusion; this relies on the physiological advantages conferred by cell-cell fusion and is independent of the question of the genetic advantages (and disadvantages) of sexual reproduction. This alternative perspective offers useful new insights that can be compared with empirical observation.

That size-based processes could play a role in the early evolution of sexual reproduction has empirical and theoretical support. The “food hypothesis” [28] suggests that metabolic uptake could drive horizontal gene transfer in bacteria and archaea, with DNA molecules providing nutrients for the receiving cell [29, 30]. Indeed, horizontal gene transfer has been shown experimentally to be an important source of carbon and nutrients in bacteria [31, 32]. Binary cell fusion is possible in bacteria (where it has been shown to come with selective benefits from mixed cytoplasm [33]) and archaea [34, 35].

Meanwhile amongst eukaryotes, the benefits of increasing cytoplasmic volume are understood to be strong enough to drive selection for the sexes themselves [36, 37]. That early selection for syngamy may have been driven by the survival benefits of larger cytoplasmic volume was argued verbally by Thomas Cavalier-Smith. He suggested that syngamy’s initially prime role was “to make zygotes larger and increase their survival rate by being able to store more solid food reserves” [38] and that this selective pressure would have been particularly important under stressful conditions “where life was threatened already by famine and the cell had less to lose and more to gain by fusing with others” [39]. However this hypothesis has until now not been explored mathematically. In suggesting a mechanistic hypothesis for the evolution of binary cell fusion, our work has interesting parallels with [40], where an advantage to cell fusion is identified in terms of shortening the cell-cycle.

Moving to consider potential physiological benefits of binary cell fusion naturally leads to work on the evolution of multicellularity. While multicellularity and binary cell fusion are clearly biologically distinct, from a modelling perspective they share similarities in that they can involve the “coming together” of cells to produce a larger complex [41]. Multicellularity achieved via aggregation allows organisms to rapidly adapt to novel environments that favour increased size [42, 43]. It has also been suggested that the genetic nonuniformity of such aggregates may also make them well-suited to resource limited environments [44, 45], echoing the hybrid vigor hypotheses for the evolution of early syngamy [4]. Relatively few theoretical studies have investigated the evolution of facultative aggregation in response to changing environments [43]. However in the context of clonal multicellularity (“staying together”) such environments have been considered more extensively [46, 47]. In this clonal context, the evolutionary dynamics act primarily on fragmentation modes [48, 49] (e.g. how a “parental” multicellular complex divides to form new progeny). Interestingly the same quality-quantity trade-off arises here [46] as drives selection for the sexes [36] (anisogamy, gametes of differing sizes); larger daughter cells (or gametes) are more able to withstand unfavourable environmental conditions, while smaller cells can be produced in larger quantities.

In this paper we adapt the classic Parker-Baker-Smith [36] (PBS [37]) model for the evolution of sexes in order to investigate the evolution of binary cell fusion. This builds on recent work that investigates how the possibility of parthenogenetic reproduction can drive selection for oogamy in eukaryotes [50]. We assume for simplicity that parental cells undergo a number of cell-divisions. The size of daughter cells is a compound evolvable trait determined both by the size of parental cells and the number of cell divisions. Daughter cells are then introduced to a pool in which they can undergo binary cell fusion, with larger fused cells experiencing a survival advantage. Unlike in the classic PBS model, the fusion rate is also a trait subject to evolution. In the following sections we proceed to outline some insights developed from numerical simulations before going on to develop analytical results for the model in a fixed environment. Finally we introduce switching environments and show that under plastic phenotypic responses, facultative binary cell fusion in response to harsh environmental conditions can evolve.

## Model

Many theories posit that the evolution of sex took place in the context of a haploid-dominant life cycle, whereby haploid unicellular organisms reproduce asexually via mitosis and the formation of a diploid cell is a transient state that follows syngamy [38, 39, 51]. This is consistent with many extant unicellular eukaryotes, with haploid dominant life cycles prevalent in fungi [52], the norm in chlorophyte and charophyte algae [53] and accounting for approximately 30% of protist life-cycles [54]. For clarity we will therefore couch our model in terms of a population of free-living haploid cells and draw examples from these extant organisms to motivate our model. However we also stress that as our model is primarily concerned with cell mass it is somewhat agnostic to the genetic details of ploidy. Thus the same model can be used in the context of theories that suggest diploidy preceded the evolution of syngamy [55], albeit with different interpretation of the content of the cell nucleus.

Among extant unicellular haploid species that feature facultative sexual reproduction, the green alga *C. reinhardtii* is a particularly useful model organism to consider, retaining as it does important features of the LECA [56]. We therefore briefly describe its life cycle here. *C. reinhardtii* is an aquatic, single-celled haploid organism [57]. Under benign conditions, cells reproduce asexually; cells grow to increase their volume more than ten fold, before undergoing a series of *n* mitoses to produce 2^*n*^ daughter cells, or zoospores. Following dissolution of the mother cell wall, these daughter cells are released in order to complete the cycle.

This vegetative mode of reproduction is discontinued under nitrogen-limited conditions [58]. In this second scenario, *C. reinhardtii* cells instead undergo differentiation to form sexually competent gamete cells (gametogenesis). Broadly, these cells are morphologically similar to vegetative cells [59] but show enhanced motility, low photosynthetic activity and mating structures that allow for cell-cell fusion [60, 61].

Fusion between cells is not indiscriminate, but rather restricted to occurring between cells of opposite mating types [62]. These two mating types (denoted plus and minus) are genetically determined at the haploid level and can be understood as ancestral versions of the sexes [63], but with no size dimorphism between the gametes (*C. reinhardtii* is isogamous). Should nitrogen levels rise, these gametes can de-differentiate into vegetatively reproducing cells [64]. However, should nitrogen levels remain low, gametes of opposite mating type are chemotactically attracted to each other and fuse to form a binucleate cell [64]. The cell nuclei then fuse to form a diploid zygote, or zygospore, that develops a thick cell wall that is resistent to environmental stress before entering a stage of dormancy. Restoration of benign environmental conditions triggers meiosis in the zygospore and the production of four haploid daughter cells capable of vegetative growth (germination).

As addressed in the introduction, our mathematical model takes inspiration from the classic PBS model for the evolution of anisogamy [36] (the production of sex cells of different sizes). However, whereas such models typically consider the binary cell fusion (fertilization) rate a fixed parameter, we here treat it as a trait subject to evolution. In doing so, our work builds on [50], where the same core model (albeit with a different biological framework) was used to investigate the evolution of anisogamy in species capable of parthenogenesis (sex cell development in the absence of fertilization). Here we begin by reframing the model in the biological context of the early evolution of sexual reproduction, before introducing the possibility of plastic phenotypic responses (which were not considered in [50]) into the model.

## Model of cell division, growth, and survival

We begin by considering a population of individual mature cells of total mass *E*. Each mature cell has a mass *M*, such that the total number of mature cells is *E/M*. The cells may then undergo *n* ≥ 0 rounds of binary fission. Each resulting daughter cell then has mass *m* = *M/*2^*n*^, while the total number of daughter cells in the population is (2^*n*^*E*)*/M*. Following this fission phase, each daughter cell is subjected to an extrinsic mass-dependent mortality, such that larger daughter cells are more likely to survive into the next growth cycle (see upper half of Fig 1). While many choices for such a mass-dependent function are possible, in this paper we focus primarily on the Vance survival function [65], defined as

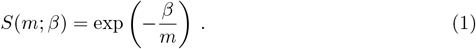

**Fig 1.**
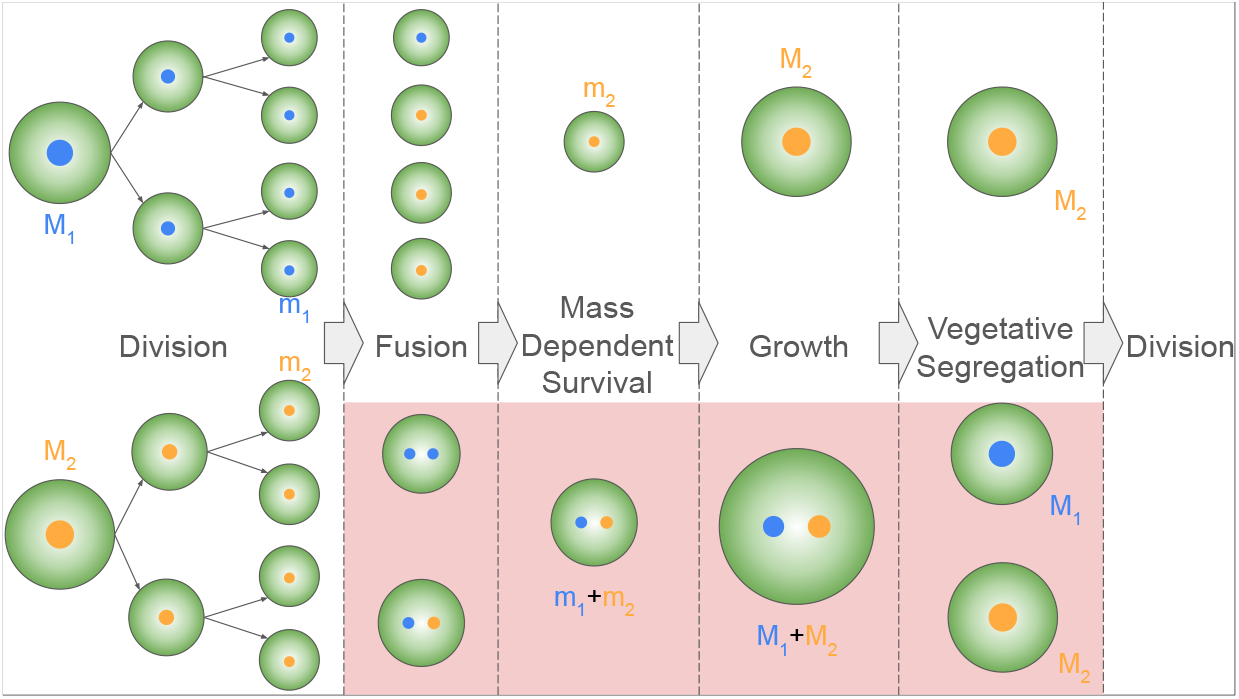
Schematic for model life cycle. Cells (green circles) are characterized by different genotypes (orange and blue nuclei) that control the non-recombining traits *m*_*i*_ and *α*_*i*_ (respectively the daughter cell mass and fusion rate of the *i*^th^ genotype). Daughter cell mass is a compound parameter that depends both on the mature cell mass *M*_*i*_ and the number of cell divisions 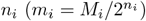 and is treated as a single continuous parameter. Following release of the maternal cell wall, daughter cells are exposed to each other. Should genotypes in the population evolve non-zero fusion rates (*α*_*i*_ *>* 0), binary cell fusion between cells is possible (see Eq. (3)). With a finite time-window for cell fusion, *T*, only a fraction of cells will be fused at the end of this fusion period. Following this both fused and unfused cells are exposed to a mass-dependent survival function (see Eq. (1)) under which larger fused cells experience a survival benefit as a result of their increased cytoplasmic volume. Surviving fused cells can grow larger due to their increased size at the beginning of the growth period. The mathematical model assumes that the size of daughter cell size of each genotype is conserved (see Eq. (4)). This comes with the implicit assumption the fused cells undergo an additional round of cell division that allows for segregation of nuclei in daughter cells. Costs to cell fusion can arise due to failed cell fusion, hindered growth of the fused cell and failed segregation, as described in the main text. These periods during which these costs can manifest are highlighted in red and captured mathematically by the parameter 1 ≥ *C* ≥ 0.

Here the parameter *β* describes the magnitude of the mortality process (i.e. the harshness of the environment); for a given value of *m*, an increase in *β* decreases the survival probability of daughter cells. We therefore refer to *β* as the environmental harshness parameter, with high *β* corresponding to harsh environments in which survival is difficult, and low *β* corresponding to more benign environments in which even cells of modest mass have a high probability of surviving. We note that Eq. (1) is both mathematically simple (being defined by only one parameter, *β*) and has the biologically reasonably property of capturing the principle of a minimum cell size (see also Figure 7) and is thus a common assumption in the literature [66–68]. Surviving cells then go on to grow to mass *M* and seed the next generation, which will again consist of *E/M* cells. In this way, *E* can be understood as setting the carrying capacity of the population in terms of its total mass.

We now wish to incorporate evolutionary dynamics to the simple picture described above. We note that the mass of daughter cells, *m* = *M/*2^*n*^, is dependent both on the mature cell size *M* and the number of cell divisions, *n*. For simplicity we therefore treat *m* as a genetically determined continuous trait, and explore its evolution using simulations. We note that this treatment is biologically equivalent to the evolving the cell sizer mechanism in *Chlamydomonas*, which operates to ensure that daughter cells have a uniform size [53].

Suppose that we have *S* genotypes in the population, such that each genotype produces daughter cells of mass *m*_*i*_ for *i* ∈ *S*. We further suppose that initially the frequency of the *i*^th^ genotype is *f*_*i*_, such that the number of mature cells of the *i*^th^ type at the beginning of a growth cycle is *f*_*i*_*E/M*_*i*_. These mature cells then produce *f*_*i*_*E/m*_*i*_ daughter cells, of which a fraction *S*(*m*_*i*_; *β*) survive to the next growth cycle.

Renormalizing by the total number of surviving cells, the frequency of type *i* cells that survive, 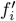 is

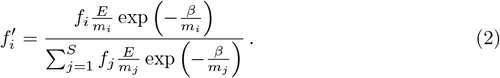

Since each of these surviving cells grows to a size *M*_*i*_, the next growth cycle begins with 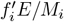 mature cells.

Note that to increase the number of daughter cells they produce (*E/m*_*i*_) mature cells can grow to smaller sizes (reduced *M*_*i*_, which increases the number of mature cells in the population) or increase their number of cell-divisions (increased *n*_*i*_). However by decreasing *M*_*i*_ and increasing *n*_*i*_, individuals also produce smaller daughter cells that are more vulnerable to extrinsic mortality. The size of daughter cells is thus subject to a quality-quantity trade-off and over multiple iterations of the deterministic cycles described above, certain genotypes may come to dominate the population. Mutants are introduced stochastically at the start of a growth cycle with average rate *µ* (the number of growth cycles between successive mutations is taken from a geometric distribution parameterized by rate *µ*). Mutant genotypes are also chosen stochastically; ancestral genotypes are chosen with probability *f*_*i*_, and mutants either increase or decrease the daughter cell mass trait of their ancestor (i.e. *m*_*S*+1_ = *m*_*i*_ *± δm*), each with an unbiased 50% probability. The parameter *δm* thus captures the size of mutational steps in trait space.

Fig 2 summarizes the outcome of such evolutionary dynamics. Fig 2A shows that the population evolves towards an evolutionarily stable strategy (ESS) in *m* for a given environment. Should the environment suddenly become harsher (via an increase in *β*) the population evolves towards a new ESS, in which daughter cells are larger (i.e. daughter cells evolve to become larger to withstand more adverse conditions).

**Fig 2.**
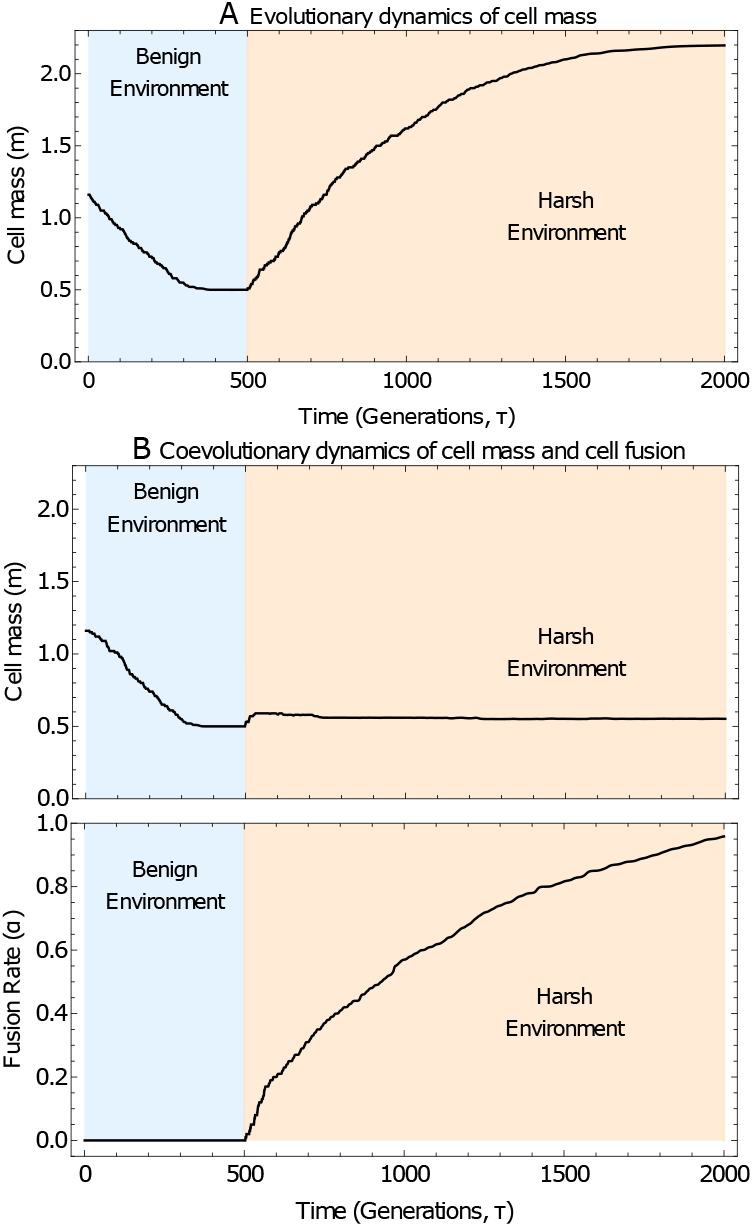
Selection for cell fusion as an alternative to increased cell size in response to a harsh environment. Stochastic simulations of evolutionary trajectories when the system is subject to a switch from the benign environment (*β*_1_ = 0.5, green region) to the harsh environment (*β*_2_ = 2.2, orange region) at growth cycle 500. Panel A illustrates the case where the fusion rate is held at *α* = 0, representing the scenario where the physiological machinery for fusion has not evolved. Panel B illustrates the case where fusion rate is also subject to evolution. Remaining model and simulation parameters are given in Supporting Information S4 and the initial condition is (*m*(0), *α*(0)) = (1.16, 0).

## Model additionally incorporating binary cell fusion

We now modify the model to allow for the possibility of binary cell fusion following the cell fission described above (see Fig 1). Daughter cells may now fuse to form a binucleated cell (e.g. a dikaryon [69], in which the cytoplasm of the contributing cells are mixed but their nuclei or nucleoids remain distinct [38]) or remain a mononucleated cell (with a single nucleus or nucleoid). The rate of cell fusion is given by *α*. We assume that this second trait is genetically linked to the daughter cell mass trait, such that each of the *S* genotypes is now defined by a trait pair (*m*_*i*_, *α*_*i*_). Cell fusions occur as a result of mass-action kinetics. For simplicity, we assume that fusions between distinct genotypes occur at their average fusion rate, (*α*_*i*_ + *α*_*j*_)*/*2. Denoting by *N*_*i*_ the number of unfused cells of genotype *i* and *F*_*ij*_ the number of fused cells resulting from daughter cells of genotype *i* and *j* we have

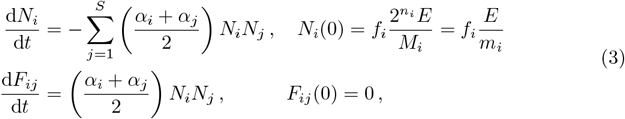

with *α*_*i*_ ≥ 0 ∀ *i*. These fusion dynamics are allowed to run for a fixed period *T*. When *α* = 0 all cells remain mononucleated. Rates *α >* 0 can be biologically understood as the evolution of any traits the promote cell fusion, such as the loss of the cell wall in the unicellular red algae *Galdieria* [70] or the formation of a mating structure in the unicellular green algae *Chlamydomonas* [71]. For *α >* 0, some proportion of daughter cells will have fused in the period *T*, leaving some cells unfused; the total number of unfused cells of type *i* is then *N*_*i*_(*T*) and the total number of fused cells with *i* and *j* nuclei are *F*_*ij*_(*T*).

We assume that cells that remain unfused at the end of the fusion period *T* can still survive and subsequently grow vegetatively. Essentially we are assuming that at early stages of this evolutionary trajectory daughter cells are not overly specialized for strict cell fusion (e.g. full gametic differentiation). This is akin to asexual reproduction in cell wall-less *Galdieria* [70] and gametic de-differentiation in *Chlamydomonas* following exposure nitrogen compounds that can be used for growth [58, 72]. For simplicity, we further assume that any potential dedifferentiation is costless. Unfused cells again survive according to Eq. (1) at a mass-dependent rate. The final number of unfused cells that survive of type *i* is therefore given by *S*(*m*_*i*_; *β*)*N*_*i*_(*T*).

Fused cells will receive a survival advantage from their increased mass, *m*_*i*_ + *m*_*j*_. As these fused cells enter the growth period with a larger inital immature cell size, we also assume that fused cells can grow to a larger mature size, *M*_*i*_ + *M*_*j*_. In order for this mature cell to produce daughter cells of the same size as unfused cells (e.g. controlled by a sizer mechanism as in *Chlamydomonas* [53]) we implicitly assume an additional round of cell division that allows for the production of mononucleated progeny through vegetative segregation [73] (or alternatively through plasmid segregation machinery [74]), as illustrated in Fig 1.

In practice, the evolution of the machinery required for cell-fusion (and subsequent segregation) is likely to come with costs. These costs include: (i) cell-fusion failure [75] in the initial fusion phase; (ii) selfish extra-genomic elements in the cytoplasm [15], cytoplasmic conflict [16, 17] and maintenance of a binucleated cell [76] hindering growth in the growth phase; (iii) the possibility of binucleated cells failing to form mononucleated progeny (i.e. failed segregation) in the vegetative segregation phase. These phases with potential costs are highlighted in red in Fig 1. We account for these costs with a single parameter 1 ≥ *C* ≥ 0. For (i) costs arising solely from cell fusion failure, (1 − *C*)*F*_*ij*_(*T*) fused cells survive to be exposed to the mass-dependent survival phase. For (ii) costs arising solely from inhibited growth, mature cells would reach a size (1 − *C*)(*M*_*i*_ + *M*_*j*_)). For (iii) costs arising solely from failed segregation, only (1 − *C*)*F*_*ij*_(*T*)*S*(*m*_*i*_ + *m*_*j*_) surviving mature cells would successfully segregate, with the remainder dying.

Analogously to Eq. (4) we can now write down the change in frequency of type *i* the frequency of type *i* cells that survive, 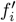 is

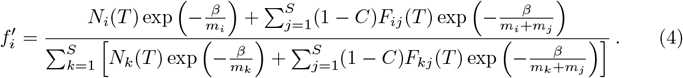

We now explore the coevolution of daughter cell mass, *m*, and fusion rate, *α*. We assume that mutations in *α* occur independently of *m* but at the same fixed rate *µ*, where *µ* is measured in units of (number of growth cycles)^−1^ (see also Supporting Information S2.5 and [50]). Analogously to mutations in daughter cell mass, ancestral genotypes are chosen proportional to their frequency, *f*_*i*_, with mutant genotypes inheriting their parental genotype plus or minus some trait deviation, *δα*; *α*_*S*+1_ = *α*_*i*_ *± δα*.

In Fig 2B, we see that in the benign environment, *α* remains at zero, and the population evolves towards an ESS in *m* as in Fig 2A. However now when the population is introduced to a harsher environment, the evolutionary dynamics differ from those in Fig 2A (where *α* was held artificially at zero). Rather than cells evolving to be larger, we see a different response emerging; selection for binary cell fusion (*α >* 0).

The result above is in some sense surprising. Despite the presence of additional survival costs associated with binary cell-fusion, selection for non-zero fusion rates (rather than increased daughter cell size) persists in the harsh environment. We explain the emergence of this behaviour mathematically in the Mathematical Analysis and Results section.

## Switching environments with phenotypic plasticity

We have seen in Fig 2B that selection for cell fusion can evolve in response to harsh environmental conditions. We are particularly interested in exploring when such cell fusion can evolve as a plastic stress response to changing environmental conditions, as observed in the sexual reproduction of unicellular organisms such as *Tetrahymena* [19, 20], *C. reinhardtii* [21] and *S. pombe* [22]. We recall that while the core model in a fixed environment (described above) has been analysed in [50], the possibility of phenotypic plasticity was not considered.

We model environmental change as switching between two environments *β*_1_ and *β*_2_. If *β*_2_ *> β*_1_, then *β*_2_ is the harsher environment in which cells have a lower survival probability (see Eq. (1)). For clarity, we will keep this convention for the remainder of the paper. We will also allow for phenotypic plasticity such that the population can evolve different strategies in different environments. The population’s evolutionary state is now described by four traits; the daughter cell mass in environments 1 and 2 (*m*_1_ and *m*_2_) and fusion rate in these environments (*α*_1_ and *α*_2_). We assume that these traits are all linked, so that recombination between the traits is not possible. For simplicity we initially assume that any cost of phenotypic switching or environmental sensing is negligible and that this plastic switching is instantaneous upon detection of the change in environmental conditions.

## Implementation of Numerical Simulations

The stochastic simulations of the evolutionary trajectories are also implemented using a Gillespie algorithm [77] where successive mutations and environmental switching events occur randomly with geometrically distributed waiting times. The rates of mutations *µ* and environmental switching *λ* are measured in units of (number of growth cycles)^−1^. In the simulations, multiple traits coexist under a mutation-selection balance (see Supporting Information and [50] and [78] for more detail), which allows us to account for variations in selection strengths in simulations of our evolutionary trajectories.

As in [50], environmental switching is modeled as a discrete stochastic telegraph process, with the time spent in each environment distributed geometrically, the discrete analogue of an exponential distribution since the time spent in each environment is measured in the number of discrete generations. The population spends an average of *τ* _1_ = 1*/λ*_1→2_ in environment 1 and *τ*_1_ = 1*/λ*_2→1_ in environment 2, where *λ*_*i*→*j*_ is the transition rate from environment *i* to *j*.

## Mathematical Analysis and Results

In order to analyse the dynamics of the model, we use tools from adaptive dynamics [79], assuming that traits are continuous and that mutations have small effect to develop equations for the evolutionary dynamics of traits *m* and *α*. We have previously derived these equations in [50] for the case without phenotypic plasticity. In [50] we were primarily concerned with the evolution of suppressed fertilization (gamete fusion) in sexual eukaryotes that had evolved the capacity for asexual reproduction. Mathematically, this equates to considering initial evolutionary conditions *α*(0) *>* 0. In contrast, in the present paper we focus on the initial evolution of cell fusion, which is equivalent to considering possible evolutionary outcomes when *α*(0) = 0.

For orientation, we begin by describing the possible evolutionary trajectories from an initial state (*m*(0), *α*(0)) = (*m*_0_, 0) in a fixed environment (i.e. when the parameter *β*, which measures the harshness of the environment (see Eq. (1)), is constant). We then move on to exploring the consequences of these dynamics for a population in switching environments with a plastic phenotypic response.

## Fixed Environment

Adopting the classical assumptions of adaptive dynamics [79, 80], we obtain equations for the selective gradient on traits *m* and *α*, which we write as *H*_*m*_(*m, α*; *β, C*) and *H*_*α*_(*m, α*; *β, C*) respectively (see Supporting Information S1 and [50]). We find that along the zero-fusion boundary, the selection gradient *H*_*α*_(*m*, 0; *β, C*) can be negative. While this makes sense mathematically (it is equivalent to the additional *production* of cells during the fusion window, see Eq. (3)) it makes no sense biologically (the population’s cell mass is not conserved over the cell division and fusion period). We therefore introduce a biologically realistic boundary at *α* = 0. The equations for the evolutionary dynamics of *m* and *α* are then given by

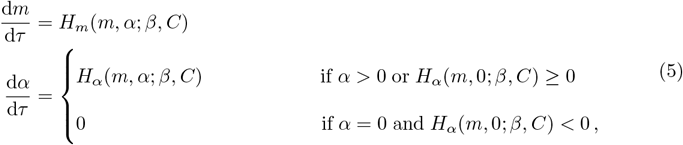

with

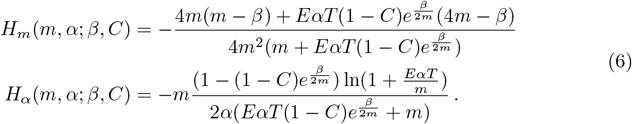

In total we find three possible attractors for the evolutionary dynamics (see Supporting Information Section S2), which we outline below.

We begin by considering the dynamics along the *α* = 0 boundary. For *α* = 0, the equation for the evolutionary dynamics of *m* simplifies to

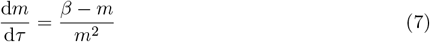

while the inequality *H*_*α*_(*m*, 0; *β, C*) *<* 0 simplifies to

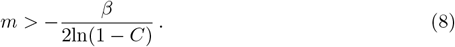

These equations, together with the condition *α* ≥ 0, suggest that a fixed point exists on the *α* = 0 boundary at

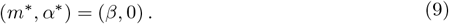

Meanwhile for large *α* we find an additional diverging trajectory. In the limit *α* → ∞, we find

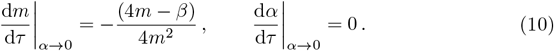

These equations suggest that an attracting manifold *m* = *β/*4 exists in the limit *α* → ∞

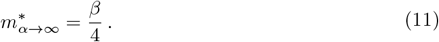

The third and final potential evolutionary attractor is found by solving *H*_*m*_(*m, α*; *β, C*) = 0 and *H*_*α*_(*m, α*; *β, C*) = 0 for arbitrary *m* and *α*. We obtain

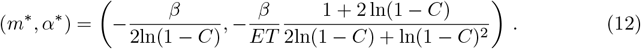

Unlike the previous evolutionary attractors (see Eqs. (9–11)), this fixed point is a function of the cost to cell-fusion *C*. It is therefore useful to focus on the behaviour of this fixed point as *C* is increased in order to characterize the behaviour of the population in different parameter regimes, as illustrated in Fig. 3.

**Fig 3.**
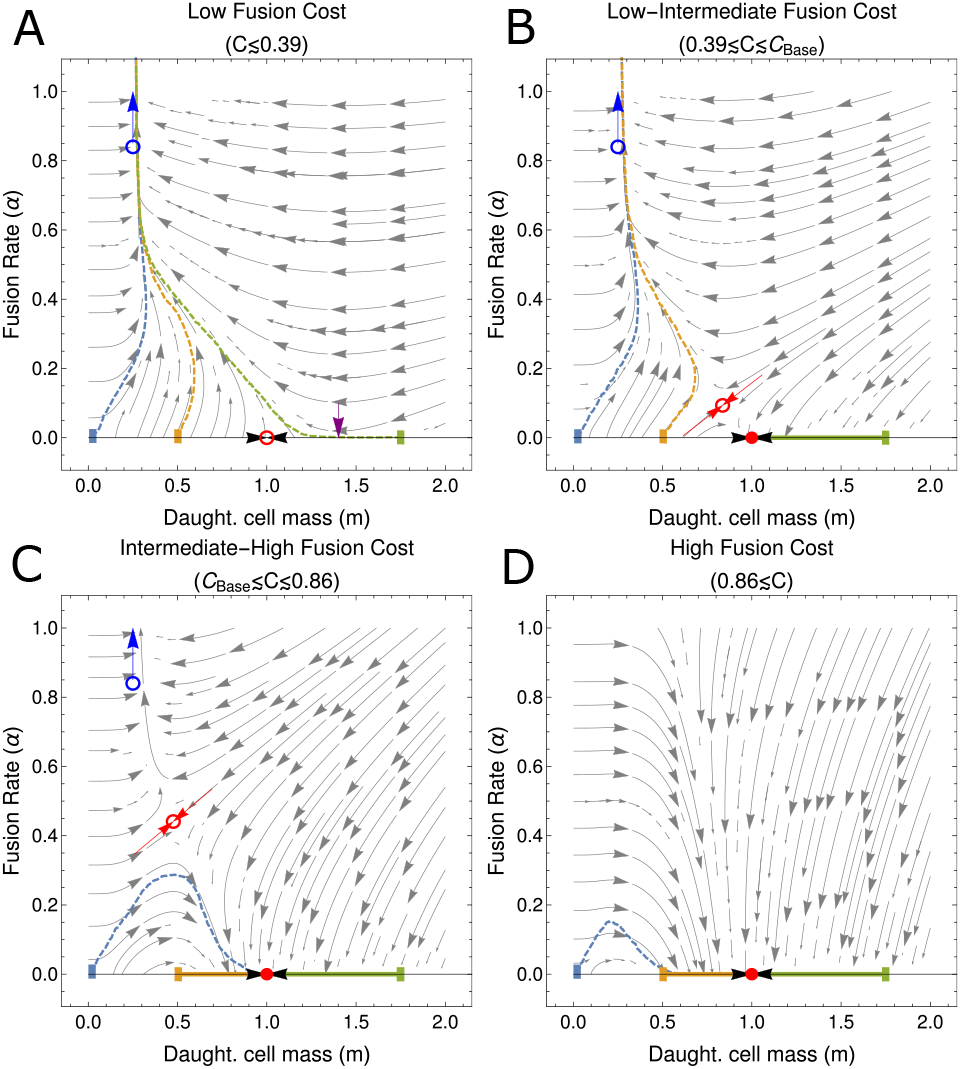
Fusion costs govern the basin of attraction for the zero-fusion state in a fixed environment. Each panel gives the phase portrait for the co-evolutionary dynamics of (*m, α*) in a fixed environment (see Eq. (5)). (A) High fusion rates are the only evolutionary outcome when costs to cell fusion are low, with an unstable fixed point at (*m*, *α*) = (*β*, 0) (open red circle) and selection for increasing *α* on the invariant manifold *m* = 1*/β* (blue arrow). Purple arrow illustrates the point on the *m* = 0 boundary when selection for *α* becomes positive (see Eq. (8)). (B) Under low-intermediate fusion costs, a third saddle fixed point emerges (see Eq. (12)), undergoing a bifurcation with (*m*, *α*) = (*β*, 0) and rendering it stable (solid red disk). There are now two evolutionary outcomes, depending on initial conditions; trajectories either increase in *α* along the invariant manifold, (*m*, *α*) → (*β/*4, *∞*), or are attracted to the (*m*, *α*) = (*β*, 0) fixed point. (C) For intermediate to high fusion costs, the separatrix of the saddle fixed point (red arrows) crosses the origin, rendering the entire *m* = 0 boundary within the basin of attraction of the (*m*, *α*) = (*β*, 0) fixed point. While for low initial cell masses the fusion rate may initially increase, ultimately the population will evolve to a low fusion state. (D) Under high fusion costs the saddle fixed point crosses the invariant manifold *m* = 1*/β*, reversing selection for the fusion rate such that d*α/*d*τ* is now negative along this line. In this scenario all initial conditions evolve towards the (*m*, *α*) = (*β*, 0) fixed point. Coloured dashed lines show the average over multiple stochastic realizations of the simulated model (see also Fig. S1) with initial conditions *m*(0) = 0.02 (blue), *m*(0) = 0.5 (orange), *m*(0) = 1.75 (green) and all in an initial state without fusion (*α*(0) = 0). Each simulated trajectory is run for 1.5 × 10^6^ growth cycles and remaining parameters are given in Supporting Information S4.

When *C* = 0, the fixed point given in Eq. (12) yields a negative, and therefore biologically nonphysical, value of *α*. The remaining biologically relevant evolutionary attractors are an unstable saddle at the fixed point along the *α* = 0 boundary (see Eq. (9)) and a diverging trajectory towards *α* → ∞ along the *m* = *β/*4 manifold (see Eq. (11)), as illustrated in Fig. 3(A). As *C* is increased, *α* at the fixed point given in Eq. (12) also increases, eventually crossing the *α* = 0 boundary when *C* = 1 − *e*^−1*/*2^ (see expressions for *m* in Eqs. (9) and (12)). At this point the fixed point given by Eq. (12) crosses the fixed point along the *α* = 0 boundary and a bifurcation occurs, yielding a saddle fixed point on the interior and rendering the fixed point at the boundary stable (see Fig. 3(B)). At this stage the basin of attraction for the stable fixed point on the *α* = 0 boundary only contains larger values of *m* for initial conditions with *α*(0) = 0. As the cost to cell fusion, *C*, becomes larger still, the basin of attraction for the stable fixed point on the *α* = 0 boundary grows to encompass the entire *α* = 0 boundary. We denote the critical value of *C* at which this occurs as *C*_Base_. At this stage, should *m*_0_ be sufficiently small, an initial increase in fusion can be observed but the population is ultimately attracted to a zero-fusion state at long times, as illustrated in Fig. 3(C). Finally as *C* is increased to very large values, the interior fixed point crosses the diverging trajectory with *α* → ∞ when *C* = 1 − *e*^−2^ (see expressions for *m* in Eqs.(11) and (12)). This results in another transcritical bifurcation, following which selection for *α* along the *m* = *β/*4 manifold becomes negative, and the fixed point given in Eq. (12) becomes biologically nonphysical (see Fig. 3(D)). This bifurcation then leaves the zero fusion state as the only stable fixed point.

The scenarios described above for *C > C*_Base_ are qualitatively similar, in that they feature a single evolutionary endpoint for all trajectories with initial conditions (*m*(0), *α*(0)) = (*m*_0_, 0) (see in Fig. 3(C-D)). In order to determine *C*_Base_ analytically, we need to obtain the value of *C* for which the separatrix, marking the boundary of the basin of attraction for the fixed point (*m*, *α*) = (*β*, 0), intersects the origin point (*m, α*) = (0, 0). We can obtain a linear approximation for this separatrix using the stable directions of the fixed interior saddle fixed point given in Eq. (12) (see red arrows in Fig. 3(B-C)). In the Supporting Information Section S2.4 we show this is given approximately by the solution to

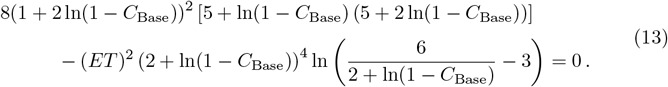

The root of this equation can be understood as being governed by the compound parameter *ET* (see Fig. 4). When *ET* is small (i.e. the density of cells entering the fusion pool is low or the period allowed for fusion short), the root approaches the solution to 1 + 2 ln(1 − *C*_Base_) = 0. Meanwhile when *ET* is large the root approaches the solution to 2 + ln(1 − *C*_Base_) = 0. These limits to the solution for *C*_Base_ are precisely the bifurcation points identified previously.

**Fig 4.**
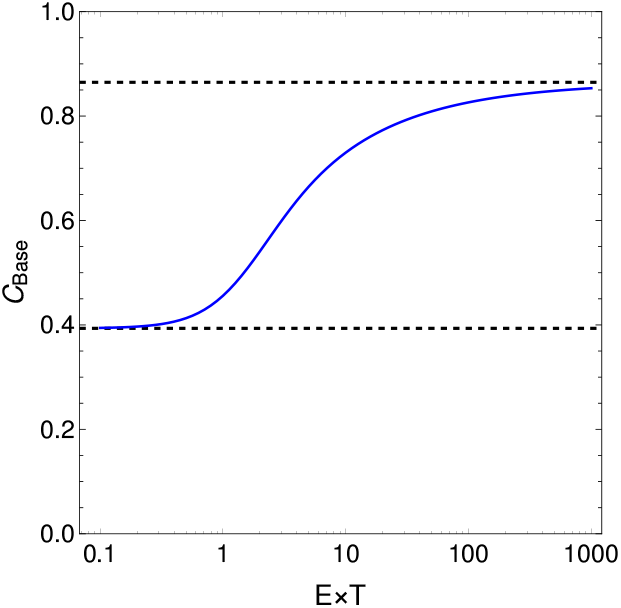
Critical cost above which cell fusion becomes impossible to evolve increases with cell density and time available for cell fusion. Blue line shows numerical solutions of Eq. (13). For values of *C > C*_Base_, obligate cell fusion remains a potential evolutionary outcome (see Eq. (14)). The value of *C > C*_Base_ is bounded above and below by the bifurcation points identified in the main text (black dashed lines, see Eq. (15)).

Summarizing the results above, we therefore have three possible evolutionary scenarios starting from initial conditions (*m*(0), *α*(0)) = (*m*_0_, 0). Which of these scenarios we observe depends on the cost to cell fusion, *C*. When *C* is low, the only evolutionary endpoint is the invariant manifold at high fusion rate. In this scenario, obligate fusion is the only evolutionary outcome, irrespective of *m*_0_. For intermediate costs to cell fusion, there are two possible evolutionary outcomes, depending on the initial daughter cell mass *m*_0_: if *m*_0_ is small, selection acts to increase fusion rate and obligate fusion is the ESS; if conversely if *m*_0_ is large, the state of no cell fusion is the ESS. Finally when costs to fusion are high selection for decreased fusion rate acts regardless of the initial value of *m*_0_. Mathematically

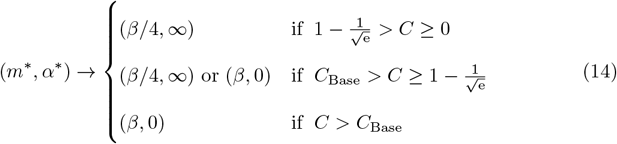

where

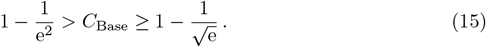

We note that it has been previously shown in [50] that the first of these conditions has additional consequences in this model for the evolution of suppressed fertilization in eukaryotic macrogametes. Should the costs to fertilization (cell fusion) exceed *C* = 1 − *e*^−1*/*2^, evolutionary branching can occur at large values of the fusion rate *α*. However this branching happens at a later evolutionary stage than the focus of this study, the early emergence of binary cell fusion.

We conclude this section by addressing the key biological result that arises from this analysis; cell fusion is uniformly selected for even under moderately high costs (with a fraction of up to *C* ≈ 0.39 of fused cells failing to survive) and can even be selected for under higher costs (up to *C* ≈ 0.86) given necessary initial conditions and parameters. In the context of the evolution of early binary cell fusion, this provides a surprising nascent advantage to cell fusion. This advantage could even help compensate for other short-term costs arising from the later evolution of sex and recombination. The selective advantage experienced by fusing cells comes from their increased cytoplasmic volume, which leads to increased survival probabilities.

## Switching environments with phenotypic plasticity

We now consider the case where evolution acts on the same traits as before, but where the environment is subject to change and the population can evolve a specific response to each environment. We find that under the assumption of costless phenotypic plasticity, the evolutionary dynamics in each environment are decoupled (see Supporting Information Section S3.2);

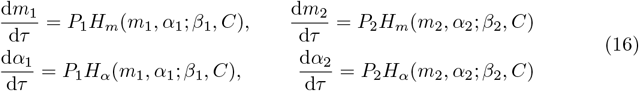

where *H*_*m*_(*m, α*; *β, C*) and *H*_*α*_(*m, α*; *β, C*) retain the functional form in Eq. (5) and we again restrict the dynamics to the biologically reasonable *α* ≥ 0. Note that the key effect of environmental switching here is to moderate the rate of evolution, with each trait pair (*m*_*i*_, *α*_*i*_) evolving at a rate proportional to the time the population is exposed to environment *i*.

While the *dynamics* of the population in each environment is decoupled, the evolutionary *trajectories* in each environment are coupled by the initial trait values for the population in each environment, which we assume are the same;

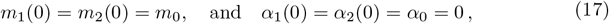

i.e. the population begins in a phenotypically undifferentiated state. As addressed, we are particularly interested in the case where cell fusion emerges as a plastic response, being selected for in one environment and against in the other. Considering the scenarios summarized in Eq. (14), this restricts our attention to the case of intermediate costs to cell fusion, *C*_Base_ *> C* ≥ 1 − *e*^−1*/*2^, for which both the zero fusion and the obligate fusion states are potential evolutionary outcomes.

As Eqs. (16) are only coupled through their shared initial conditions, *m*_0_ and *α*_0_, the choice of these initial conditions is an important consideration. Since we are interested in the initial evolution of binary cell fusion, it is natural to assume that the population evolves from a state of zero fusion, *α*_0_ = 0. Deciding on a plausible initial daughter cell mass, *m*_0_, takes more thought. We will consider two parsimonious choices.

One parsimonious choice would be that the population is already adapted to either environment 1 or environment 2 and that *m*_0_ is given by an evolutionary fixed point in one of these environments (this is the situation illustrated in Fig. 2). Suppose that the population was first exposed to environment 1 (with *β* = *β*_1_), and that in this environment there was no selective pressure to evolve cell fusion (see Eq. (9)). The population would then initially reside at the zero-fusion fixed point of environment 1,

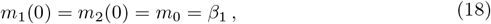

as illustrated by the purple circles in Fig. 5). In order for selection for cell fusion to act in environment 2, *m*_0_ = *β*_1_ must lie outside the basin of attraction of the stable zero-fusion fixed point *m* = *β*_2_ in environment 2. In general, the conditions required for this scenario to hold must be obtained numerically. However when *ET* is large (e.g. in high-density environments), the condition simplifies analytically to

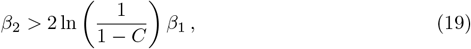

i.e. the second environment must be sufficiently harsher than the first environment in order for obligate cell fusion to evolve as a stress response. This is illustrated in Fig. 6(A).

**Fig 5.**
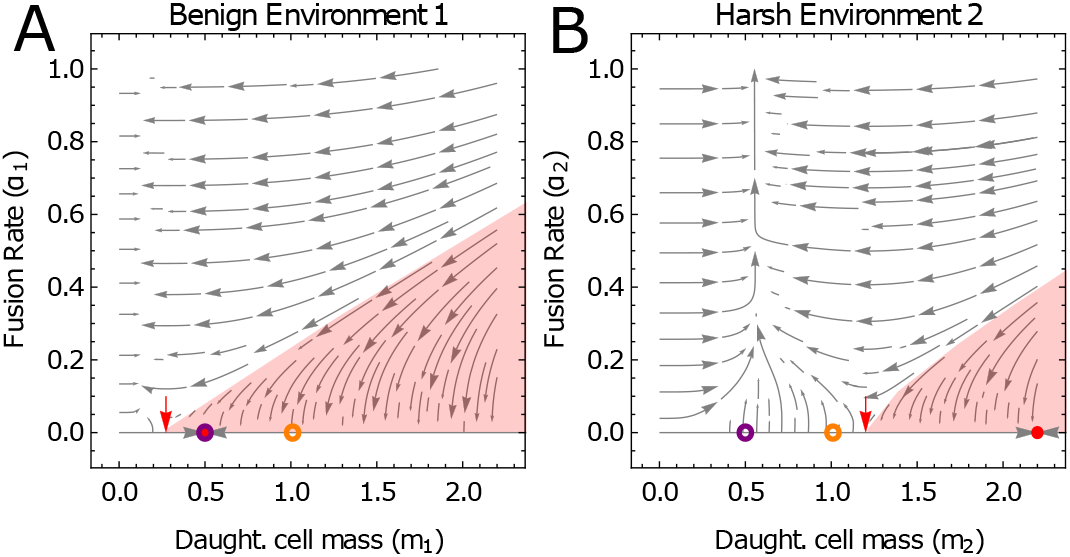
Selection for cell fusion as an alternative to increased cell size in response to a harsh environment. Illustrative phase portrait for co-evolutionary dynamics of (*m*_1_, *α*_1_, *m*_2_, *α*_2_) in a switching environment with phenotypic switching that exhibits facultative binary cell fusion. In both environment 1 (panel A) and environment 2 (panel B) the cost to cell fusion is *C* = 0.6, purple circles represent the initial condition (*m*_1_(0), *α*_1_(0)) = (*m*_2_(0), *α*_2_(0)) = (*β*_1_, 0), and orange circles represent the initial condition 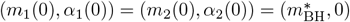 (see Eq. (20)). The red shaded regions shows the approximate region of the basin of attraction for the stable zero fusion fixed points (*m*, *α*) = (*β*_1_, 0) (see red disk, panel (A)) and (*m*, *α*) = (*β*_2_, 0) (see red disk, panel (B)). Red arrows indicate the intersection of these basins with the *α* = 0 boundary. Environmental parameters are *β*_1_ = 0.5 and *β*_2_ = 2.2 making environment 1 the more benign environment, in which the population typically spends a proportion *P*_1_ = 0.7 of its time.

**Fig 6.**
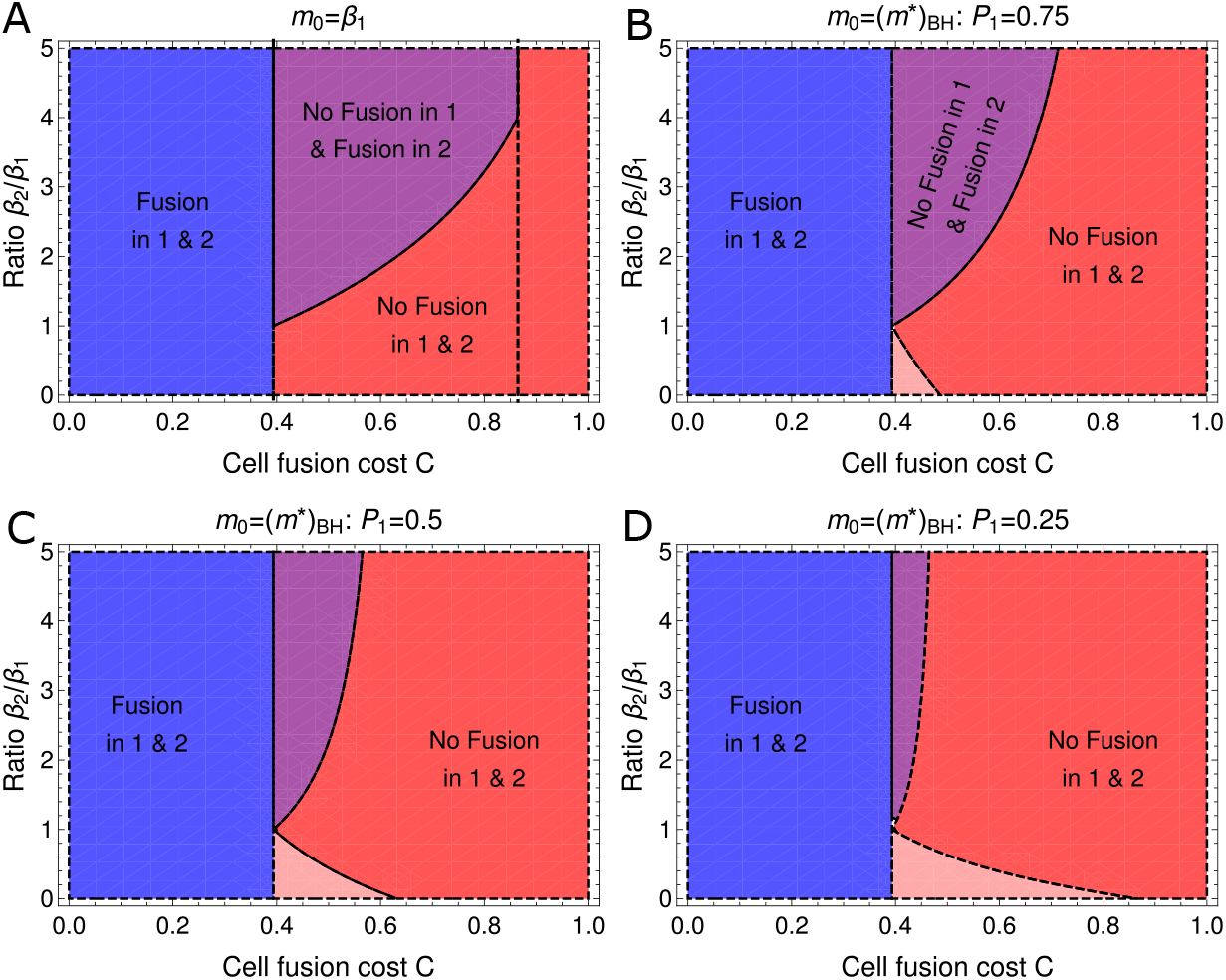
Conditions on cell fusion cost and deterioration in environment required to observe cell fusion as a facultative stress response. Regions are calculated in the limit of large *ET* and show when fusion is selected for in both environments (blue, 1 − *e*^−1*/*2^ *> C*) and in neither environment (red, e.g. *C >* 1 − *e*^−2^ *> C*) and when fusion is selected for in only the harsher environment (purple and pink). On the interval 1 − *e*^−2^ *> C >* 1 − *e*^−1*/*2^ the evolutionary outcome depends on the ratio of environmental parameters and on the initial evolutionary conditions. In panel (A) the population is initially adapted to environment 1, and the condition for facultative cell fusion in environment 2 is given in Eq. (19). In panels (B-D) the population has initially adapted a bet-hedging strategy to the two environments. Cell fusion remains at zero in environment 1 but evolves in environment 2 if Eq. (21) hold but Eq. (22) is violated (see purple regions). Conversely cell fusion evolves in environment 1 but is held at zero in environment 2 if Eq. (21) is violated but Eq. (22) holds (see pink regions).

**Fig 7.**
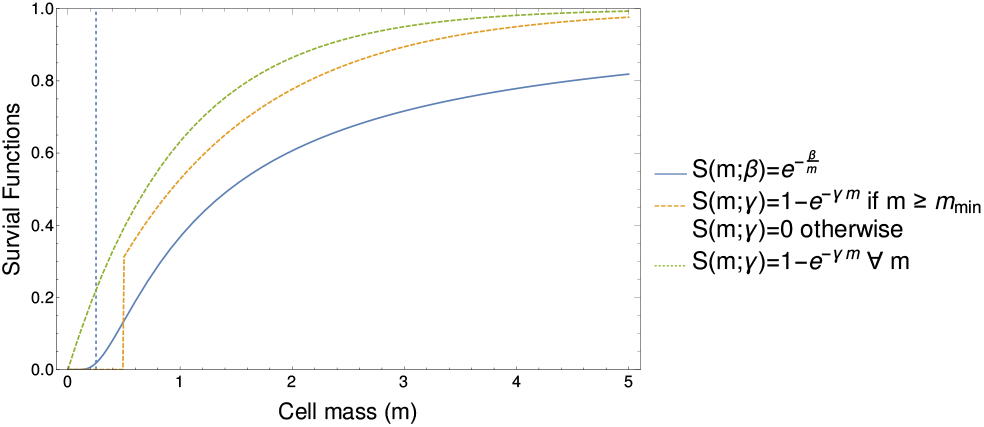
Various choices for the survival function. The blue line illustrates the Vance survival function given in Eq. (1) for *β* = 1. We note that this function implicitly captures the feature of a minimum viable cell size as which the survival of cells drops precipitously (see vertical dashed blue line at *m* = *β/*4). The green dashed line shows the survival function given in Eq. (23) with *γ* = 1 and *m*_min_ = 0. Here cell survival decays linearly with mass for small *m*. The orange dashed line shows Eq. (23) with *γ* = 0.75 and *m*_min_ = 0.5.

An alternative choice for the initial cell mass assumes that the population has evolved under exposure to both environment 1 and environment 2, but has not yet evolved phenotypically plastic responses. We have shown in [50] (see also Supporting Information S3.1) that for *α* = 0, the ESS for daughter cell cell mass in such a scenario is a bet-hedging strategy;

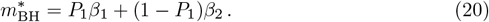

Note that this is intuitively equivalent to the *α* = 0 evolutionary strategy in environment 1 (weighted by the probability of being in environment 1) plus the evolutionary strategy in environment 2 (weighted by the probability of being in environment 2), as illustrated by the orange circles in Fig. 5). We have shown in [50] that Eq. (20) is an accurate approximation if the timescale of environmental switching is faster than the evolutionary timescale (*λ*_*i*→*j*_ *> µ*).

We must now determine whether the initial condition *m* lies in the basins of attraction of (*m*, *α*) = (*β*_1_, 0) in environment 1 and (*m*, *α*) = (*β*_2_, 0) in environment 2. Again this in general requires a numerical approach. However we show in Supporting Information Section S3.2.2 that analytic progress can be made in the limit of large *ET*. We find that in the cost region in which a facultative response is possible (1 − *e*^−2^ *> C* ≥ 1 − *e*^−1*/*2^), (*m*, 0) is within the basin of attraction of (*m*, *α*) = (*β*_1_, 0) in environment 1 if

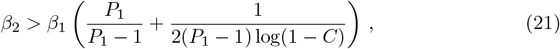

and 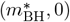 is within the basin of attraction of (*m*, *α*) = (*β*_2_, 0) in environment 2 if

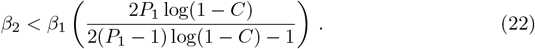

If only one of these conditions holds, then facultative cell fusion can be observed in one environment but not the other. Further, if facultative cell fusion does occur, it will always occur in response to the harsher of the two environments.

In Fig. S7(B-D) we illustrate how the possible evolutionary outcomes vary as *P*_1_ = *τ*_1_*/*(*τ*_1_ + *τ*_2_) (the probability of being in environment 1) is reduced (note that the plot for *P*_1_ → 1 tends to Fig. S7(A), see Eq. (20)). We see that as the time spent in environment 1 is reduced, so too is the parameter regime in which one would expect to see facultative cell fusion in response to a harsher environment 2. However we also see the emergence of a region in which environment 2 is more benign (*β*_2_ *< β*_1_) and in which by symmetry facultative cell fusion can emerge in response to the harsher environment 1 (see pink regions in Fig. S7(B-D)). In Figs. S5-S7 of the Supporting Information, we show that these analytic results are good predictors of the dynamics obtained via numerical simulation.

In summary, for both scenarios described above, we see how facultative binary cell fusion can evolve as a response to harsh environmental conditions that lower the survival probability of daughter cells. Such a response is possible if costs to cell fusion are intermediate and if there is an appreciable increase in environmental harshness, *β*, between the environments. We have also seen that regular exposure to both environments before the evolution of cell fusion has taken place (such that the population has previously adapted a bet-hedging strategy in daughter cells mass) may reduce the parameter region over which we expect to see cell fusion as a stress response. However we note that in many sexually reproducing unicellular eukaryotes, the environmental conditions that trigger sexual reproduction (and cell fusion during plasmogamy) are very rare; sexual reproduction is estimated to be triggered approximately once in every 2000 generations in *Saccharomyces paradoxus* and once in every 770 generations in *C. reinhardtii* [81, 82]. These values would be equivalent to *P*_1_ ≈ 1 in our model, which is the regime in which facultative binary cell fusion occupies the broadest region of parameter space.

## Alternative Survival Functions: The importance of minimum cell size

Throughout the mathematical analysis above, we have considered one specific form of survival function, namely the Vance survival function Eq. (1). In order to illustrate the robustness of our results, we explore the effect of alternative survival functions in Supporting Information Section S5. These functions are illustrated in Fig. 7. We show that an important feature of the Vance survival function is that it implicitly accounts for very small cells having a low survival probability (the growth of Eq. (1) is sub-linear when *m < β/*2 as Eq. (1) is convex in this region). Conversely the function 1 − exp(−*γm*) is linear in *m* when *m* is small. Consequently we find that the evolutionary dynamics generated by the survival function 1 − exp(−*γm*) lead to selection for the production of infinitely many (and infinitely small) daughter cells. In order to correct for this behavior, we can modify the function to account for a minimum viable cell size;

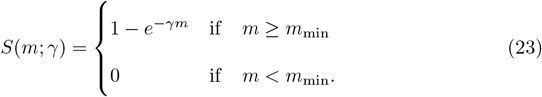

The incorporation of this minimum cell size allows us to identify analogous parameter regimes as identified when using the Vance survival function. In particular we once again find that binary cell fusion can be selected for as a stress response should one environment be sufficiently harsher. The exact parameters at which this occurs are dependent on the choice of *m*_min_ and the fusion cost, *C*. However one notable point of departure is the maximum value for *C* under which cell fusion can be selections for. Whereas we have shown that the Vance survival function can allow selection for cell fusion up to *C* ≈ 0.86, using Eq. 23 leads to a maximum cell fusion cost of *C* = 0.5.

## Discussion

The evolution of sexual reproduction and its consequences for the subsequent evolutionary trajectory of populations is of general importance in biology [7, 13, 83]. In this paper we have illustrated a reversal of the classic two fold cost of sex that appears in organisms with distinct sexes [84]; in unisexual [85], unicellular organisms, binary cell fusion can be selected for, even in the presence of substantial costs, due to a survival benefit that comes from increased mass. These results allow us to quantitatively assess the verbal hypothesis that syngamy evolved allow cells to store more food reserves and thus increase their survival rate [38]. It is particularly interesting that the benefits conferred to cell fusion through increased cytoplasmic mass are sufficient to withstand remarkably high costs; “obligate sexuality” is the *only* evolutionary outcome with costs equivalent to a loss of ~ 39% cells that attempt to fuse, and remains a potential outcome with costs of up to ~ 86% of fused cells dying.

Perhaps most interesting is the case of switching environments with phenotypic plasticity. Here we find under a broad set of biologically reasonable conditions (costs to cell fusion equivalent to 39% − 86% additional mortality to fused cells and at least moderate changes in environmental quality) that high fusion rates are selected for in harsh environments and zero fusion rates are maintained in benign environments. This behaviour parsimoniously recapitulates the empirically observed reproductive strategies of numerous facultatively sexual species, including *C. reinhardtii* [21] and *S. pombe* [22]. This mechanism, under which cell fusion evolves to increase the survival probability of daughter cells, provides a complementary perspective on the frequent evolution of survival structures (resistant to environmental stress) that form following the formation of a zygote. These include ascospores in fungi [86] and zygote-specific stress-resistant stress wall in *C. reinhardtii* [87]. Note that such correlations between sexual reproduction and the formation of survival structures are not as easily explained under genetic explanations for the evolution of sexual reproduction, where engaging in both behaviour at once constitutes a simultaneous (and therefore potentially costly) change in genotype and temporal dislocation in environment [88, 89].

The results above are particularly interesting in the case of the evolution of early binary cell-fusion as a first step in the evolution of sexual reproduction. While most studies focus on the genetic benefits of cell-fusion [90] (including a functionally-diploid dikaryotic cell [4]), or the genetic benefits of mixed cytoplasm [13, 14] (which can also come with costs [15–17, 41, 91–93]), the mechanism at play here is purely physiological. Yet, as addressed above, it naturally captures the empirical observation of binary cell-fusion as response to challenging environmental conditions, a feature absent in these earlier models. While the mechanism does not explain the evolution of sexual reproduction and genetic recombination itself, it does provide a nascent advantage to binary cell fusion that sets the stage for the evolution of sex by bringing nuclei from different cells into contact for prolonged periods. The mechanism also shows the potential to counter short-term costs associated with the initial formation of a binucleated cell. In this way the mechanism could facilitate the transition from horizontal gene transfer [94, 95] to meiotic recombination, which is advantageous when genome sizes increase in length [8]. Conceivably, if genetic recombination is beneficial for myriad genetic reasons in the long-term [8, 96], it would seem natural that it would be instigated when the opportunity arises (i.e. when physiological survival mechanisms bring nuclei into close contact). We note that it is obviously possible that the first diploid cells arose by errors in endomitosis [39, 51, 55] (essentially doubling the chromosome number within a single cell) and that meiosis first evolved in this context. Such a sequence of events is still compatible with our very general model, which can alleviate short-term costs of sex such as the energy involved in finding a partner and undergoing fusion [84]. In either scenario sexual reproduction may not be only a direct response to environmental variability [97, 98], but also to the correlated formation of a survival structure.

More generally, it is interesting to note that the conditions for facultative sexuality (e.g. harsh environmental conditions) broadly coincide with those for facultative multicellularity in both bacteria and eukaryotes, with starvation triggering the formation of fruiting bodies in myxobacteria [99, 100] and flocking in yeast [101, 102]. Meanwhile in *C. reinhardtii*, the formation of multicellular palmelloids and aggregates are an alternate stress response to sexual reproduction [103], as are the formation of fruiting bodies in *D. discoideum* [104]. In this multicellular context, the sexual behavior of *D. discoideum* is particularly interesting, as once formed, the zygote attracts hundreds of neighboring cells that are then cannibalized for the provision of a macrocyst [105]. Conceptually, these various survival strategies are reminiscent of the mechanism in our model of the evolution of binary cell fusion in response to challenging environments. However it would be particularly interesting to extend our model to the multiple-fusion case of *D. discoideum*, where many cells forgo reproductive output to contribute to the macrocyst [106], to see under what conditions the results he have obtained here still qualitatively hold.

One element absent from our model is the fusion of multiple cells, which is likely to be selected for under the assumptions implicit in our model. There would clearly be an upper-limit on the number of fusions selected for, arising from the likely multiplicative effect of the fusion cost *C*. However in this context, it is interesting to note that one of the hypotheses for the evolution of self-incompatible mating types is as a signal to prevent the formation of polyploid cells [107]. Such a mechanism could also prevent the formation of trikaryotic cells should the cost of multiple fusions be too great. Extending our modeling in this direction may help explain the second stage in models for the evolution of eukaryotic sex, the regulation of cell–cell fusion [1].

The selection pressure for binary cell fusion is dependent on the formation of a binucleated cell with an increased survival benefit arising from its larger size. Although it is reasonable to assume that a single mononuclear cell has lower total resource requirements than a multicellular complex of the same size [108], we have not considered the detailed energetics of the maintenance of two nuclei [9]. In these respects incorporating dynamic energy budget theory into the model would be an important next step [109] as it would provide a clear distinction between the survival benefits of fused cells and unfused bicellular complexes. Within our modelling framework, these two structures are broadly similar [41]. However as we have shown, increased cell-cell attraction can be selected for even in the presence of large costs that one might expect under binary cell fusion but not associate with the formation of a bicellular complex.

We have assumed for simplicity a simple cell division scheme; parental cells undergo *n* rounds of symmetric division to produce 2^*n*^ daughter cells. In the context of multicellularity, switching environments have been shown to promote binary fragmentation [46]. However non-symmetric modes can be selected for [48] reflecting the diverse modes of facultative multicellular life cycles observed in bacteria [110]. It would be interesting to incorporate our results into models of cell division that account explicitly for growth [111, 112] to determine how these results for multicellular organisms carry over to the unicellular scenario, and further how they may affect those we have shown here.

Finally, we have not explicitly modelled any sources of cytoplasmic or genetic conflict [113], which we have for simplicity included in the fusion cost *C*. Nevertheless, social conflict does emerge in this model. In a recent paper we have shown how evolutionary branching can arise, with some individuals producing fewer larger cells and others producing more numerous but smaller daughter cells [50]. This branching is driven by the same evolutionary forces that drive selection for anisogamy [114], in which context this can be viewed as sexual conflict [115]. That social conflict should arise in the formation of multicellular aggregates is well understood [41, 100, 116]. However these models typically assume cells of fixed size [47, 117]. Combining the insights derived from the evolution of anisogamy literature with the theory developed in the multicellularity literature represents another promising research direction.

As addressed above, trade-offs between cell fusion rate and mass [118], cell-energetics, inbreeding, and the possibility of multiple cell-fusion events offer interesting avenues to extend this analysis. In addition, we have not accounted for the discrete nature of divisions leading to daughter cells, costs to phenotypic switching, non-local trait mutations, or pre-existing mating types. More generally, extending our mathematical approach leveraging adaptive dynamics to switching environments in other facultatively sexual populations might prove particularly fruitful [62, 119].

In this paper we have adapted the classic PBS model [36] in two key ways; allowing the fusion rate to evolve and subjecting the population to switching environments. In doing so, we have shown its capacity to parsimoniously capture the evolution of obligate binary cell fusion, obligate binary cell fission, and stressed induced binary cell fusion in unicellular organisms. These results offer particularly interesting implications for the evolution of binary cell-fusion as a precursor to sexual reproduction, as well as suggesting common mechanistic links between the evolution of binary cell fusion and multicellularity. Moreover, our analysis emphasises the importance of exploring the coevolutionary dynamics of a range of evolutionary parameters, and of developing computational and mathematical approaches to elucidate facultative sexual reproduction.

## Supporting information

S1 Text. Supporting information.pdf

## Acknowledgments

This work has made use of Viking high performance computing service at the University of York.

## Author Contributions

**Conceptualization:** Jonathan Pitchford, George Constable

**Methodology:** Xiaoyuan Liu, George Constable, Jonathan Pitchford

**Formal Analysis:** Xiaoyuan Liu, George Constable

**Software:** Xiaoyuan Liu

**Investigation:** Xiaoyuan Liu

**Supervision:** George Constable, Jonathan Pitchford

**Writing-Original draft preparation:** Xiaoyuan Liu, George Constable

**Writing-review & editing:** Jonathan Pitchford, George Constable

