## Supporting information for "Cell size and selection for stress-induced cell fusion in unicellular eukaryotes"

**S4 Parameters for simulations 16**

**S5 Alternative Survival functions 17**

### S1 Derivation of Evolutionary Dynamics: Fixed environment

In the beginning of each growth cycle, we assume a fixed energy budget $E$ for growth and daughter cell production via binary fission. These daughter cells are then subject to a period $T$ in which they undergo cell fusion (also known as a growth cycle).

Since our model involves the coevolution of two traits, the evolutionary dynamics can also be derived using the standard approach to adaptive dynamics to derive equations for the evolutionary dynamics (see, for instance, [1, 2, 3]). The key assumptions are that the stepsize of mutations is small (so that each trait can be modelled by a continuously evolving variable) and the mutation rate is low (such that each mutation can establish in a population or be driven to extinction before the arrival of the next mutant). This allows us to simplify our analysis in the first instance to just considering a population consisting of two types, mutants and residents (we recall that we assume traits $(m, \alpha)$ are non-recombining).

We will denote mutant quantities with a hat notation. Mutants can modify the size of daughter cells that they produce following binary cell fusion ($\hat{m} = m \pm \delta m$) and the fusion rate between daughter cells ($\hat{\alpha} = \alpha \pm \delta\alpha$) relative to a resident population with traits $(m, \alpha)$. We will further denote the frequency of mutants in this population by $\hat{f}$. The fusion kinetics given in Eq. (3) of the main text then simplify to

$$\begin{aligned}\frac{\mathrm{d}N}{\mathrm{d}t} &= -\alpha N^2 - \left(\frac{\alpha + \hat{\alpha}}{2}\right) N\hat{N}\,, \quad N(0) = (1-\hat{f})\frac{E}{m} \\ \frac{\mathrm{d}\hat{N}}{\mathrm{d}t} &= -\left(\frac{\alpha + \hat{\alpha}}{2}\right) N\hat{N} - \alpha\hat{N}^2\,, \quad \hat{N}(0) = \hat{f}\frac{E}{\hat{m}}\,,\end{aligned} \tag{S1}$$

for the dynamics of unfused cells and

$$\begin{aligned}\frac{\mathrm{d}F_N^N}{\mathrm{d}t} &= \frac{1}{2}\alpha N^2; \qquad F_N^N(0) = 0 \\ \frac{\mathrm{d}F_N^{\hat{N}}}{\mathrm{d}t} &= \alpha N\hat{N}; \qquad F_N^{\hat{N}}(0) = 0 \\ \frac{\mathrm{d}F_{\hat{N}}^{\hat{N}}}{\mathrm{d}t} &= \frac{1}{2}\alpha\hat{N}^2; \qquad F_{\hat{N}}^{\hat{N}}(0) = 0\,,\end{aligned} \tag{S2}$$

for the dynamics of fused cells.

The absolute finesses of resident and mutant cells, $w$ and $\hat{w}$, is given by the absolute number of surviving cells of each type at the end of a growth cycle with a fusion window of time $T$;

$$\begin{aligned}w &= S(m;\beta)N(T) + (1-C)\left[2S(2m;\beta)F_N^N(T)\right. \\ &\qquad \left.+S(\beta, m+\hat{m})F_N^{\hat{N}}(T)\right]\,, \\ \hat{w}_m &= S(\hat{m};\beta)\hat{N}(T) + (1-C)\left[2S(2\hat{m};\beta)F_{\hat{N}}^{\hat{N}}(T)\right. \\ &\qquad \left.+S(m+\hat{m};\beta)F_N^{\hat{N}}(T)\right]\,.\end{aligned} \tag{S3}$$

Obtaining an analytic expression for the absolute fitness of the respective types therefore requires an analytic expression for the solution to Eqs. (S1-S2). Such a solution is only obtainable for all values of the mutant frequency $\hat{f}$ if the mutant differs from the resident in only one of the traits i.e. $(\hat{m}, \hat{\alpha}) = (m, \hat{\alpha})$ or $(\hat{m}, \hat{\alpha}) = (\hat{m}, \alpha)$, see [4]. Further progress can be made if we restrict ourselves to the case of small $\hat{f}$ (i.e. a small initial mutant frequency as would be seen at the beginning of an invasion).

When mutants are rare, mutant-mutant interactions and the effect that the mutant has on the resident population dynamics (captured by the term $((\alpha+\alpha')/2)N\hat{N}$ in $dN/dt$ in Eq. (S1)) can be neglected, which significantly simplifies the analysis [5, 6]. Under these assumptions, the fusion kinetics are given by

$$\begin{aligned} \frac{dN}{dt} &= -\alpha N^2 - \frac{(\alpha+\alpha')}{2}N\hat{N} \\ \frac{d\hat{N}}{dt} &= -\frac{(\alpha+\alpha')}{2}N\hat{N}\,. \end{aligned} \tag{S4}$$

The solution to these equations is given by

$$\begin{aligned} N(T) &= \frac{N_0^2\alpha T}{1+N_0\alpha T} \\ \hat{N}(T) &= \hat{N}_0 - \frac{\hat{N}_0\left(\frac{1}{N_0}\right)^{\frac{1}{2}\left(\frac{\hat{\alpha}}{\alpha}-1\right)}\left(\frac{1}{N_0}+\alpha T\right)^{\frac{1}{2}\left(1-\frac{\hat{\alpha}}{\alpha}\right)}}{1+N_0\alpha T}\,, \end{aligned} \tag{S5}$$

with corresponding solutions for the number of cells involved in cell fusions obtainable via

$$\begin{aligned} F_N^N(T) &= (N_0 - N(T))/2 \\ F_N^{\hat{N}}(T) &= (\hat{N}_0 - \hat{N}(T)) \end{aligned} \tag{S6}$$

and initial conditions $N_0 = N(0) = (1-\hat{f})E/m$ and $\hat{N}_0 = \hat{N}(0) = \hat{f}E/\hat{m}$ that are functions of traits $m$ and $\hat{m}$.

When mutants are rare (i.e. $\hat{f}$ is small) the relative fitness of the mutant, $\hat{W}$ can be approximated by

$$\hat{W} = \left[\frac{\mathrm{d}}{\mathrm{d}\hat{f}}\left(\frac{\hat{w}}{w+\hat{w}}\right)\right]\Bigg|_{\hat{f}=0} \tag{S7}$$

$$\begin{aligned} &= -\frac{mEe^{\beta\left(\frac{1}{m}-\frac{1}{m+\hat{m}}-\frac{1}{\hat{m}}\right)}(m+\alpha ET)^{1-\frac{\alpha+\hat{\alpha}}{2\alpha}}}{\hat{m}\left(\alpha ET(C-1)e^{\frac{\beta}{2m}}-m\right)} \\ &\quad\times E^{\frac{\alpha+\hat{\alpha}}{2\alpha}}\left[(C-1)e^{\frac{\beta}{\hat{m}}}\left(m^{\frac{\alpha+\hat{\alpha}}{2\alpha}}-(m+\alpha ET)^{\frac{\alpha+\hat{\alpha}}{2\alpha}}\right)+e^{\frac{\beta}{m+\hat{m}}}m^{\frac{\alpha+\hat{\alpha}}{2\alpha}}\right]. \end{aligned} \tag{S8}$$

Substituting in $\hat{m} = m + \delta m$ and $\hat{\alpha} = \alpha + \delta\alpha$, we obtain the evolutionary dynamics from the fitness gradient as

$$\nabla\hat{W} = \left(\frac{\mathrm{d}\hat{W}}{\mathrm{d}\delta m}, \frac{\mathrm{d}\hat{W}}{\mathrm{d}\delta\alpha}\right)\Bigg|_{\delta m=\delta\alpha=0} = (H_m(m,\alpha;\beta,C), H_\alpha(m,\alpha;\beta,C)) \tag{S9}$$

with

$$\begin{aligned} H_m(m,\alpha;\beta,C) &= -\frac{4m(m-\beta)+\alpha ET(1-C)e^{\frac{\beta}{2m}}(4m-\beta)}{4m^2(m+\alpha ET(1-C)e^{\frac{\beta}{2m}})} \\ H_\alpha(m,\alpha;\beta,C) &= -m\frac{(1-(1-C)e^{\frac{\beta}{2m}})\ln(1+\frac{\alpha ET}{m})}{2\alpha(\alpha ET(1-C)e^{\frac{\beta}{2m}}+m)}\,, \end{aligned} \tag{S10}$$

as given in the main text. We note that identical equations are obtainable should one use alternative approximation techniques that rely on calculating the dynamics of a mutant during an invasion [4].

As described in the main text, when evaluating Eq. (S10) on the $\alpha = 0$ boundary, we find that conditions exist under which evolution selects for negative $\alpha$

($H_\alpha(0, \alpha; \beta, C) < 0$). While negative values of the fusion rate $\alpha$ can make sense mathematically (see Eq. (S1)) they are clearly unrealistic biologically. We therefore impose the artificial boundary $\alpha = 0$ in our evolutionary simulations and mathematical analysis. The resulting evolutionary equations are then in the form of a discontinuous dynamical system [7].

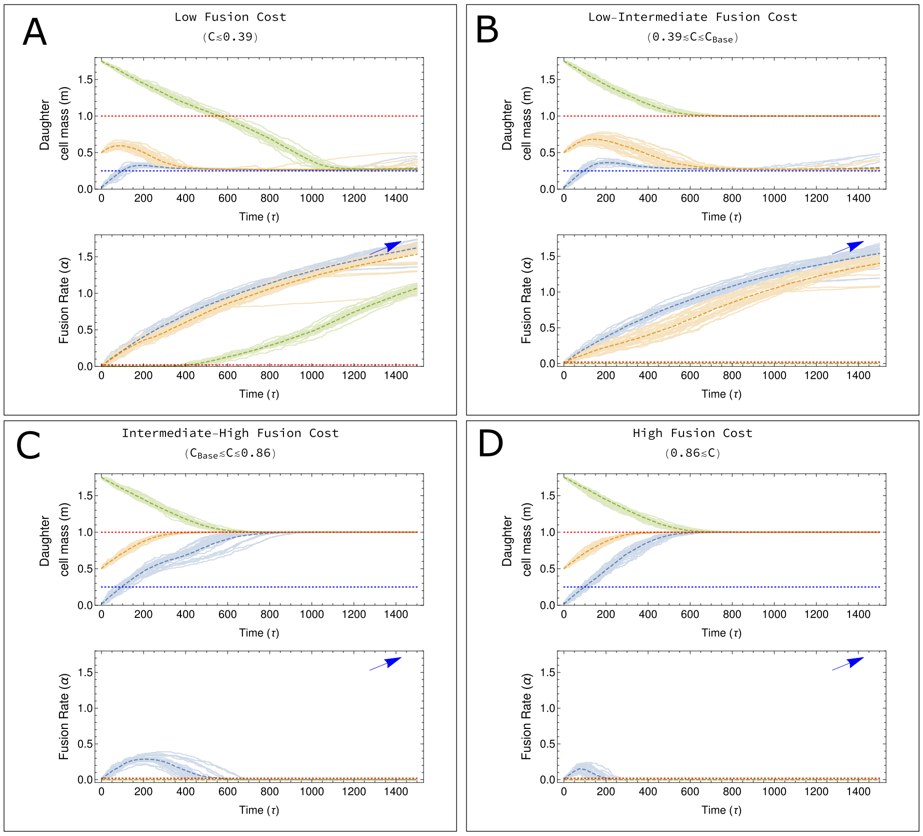

**Fig S1. Temporal trajectories for the co-evolutionary dynamics in a fixed environment.** Each panel corresponds to the trajectories plotted in Fig. 3 of the main text. Light blue, orange and green lines show the results of stochastic simulations of the model and the corresponding dashed lines show the average of these stochastic trajectories. At long times the population either tends to $(m, \alpha) = (\beta, 0)$ (see horizontal red dotted line) or $(m, \alpha) = (\beta/4, \infty)$ (see horizontal blue dotted line in plots for $m(\tau)$ and blue arrow in plots for $\alpha(\tau)$). For trajectories approaching very high values of $\alpha$, departures from the mean can be observed (see panels (A) and (B)). These departures are the result of evolutionary branching at fusion rates, as described in [4].

### S2 Analysis of Evolutionary ODEs: Fixed Environment

In the main text we identify three potential evolutionary endpoints for the dynamics, given by

$$
\begin{aligned}
&(m^*, \alpha^*) = (\beta, 0) \\
&(m^*, \alpha^*) \to (\beta/4, \infty) \\
&(m^*, \alpha^*) = \left( -\frac{\beta}{2\ln(1-C)}, -\frac{\beta}{ET} \frac{1 + 2\ln(1-C)}{2\ln(1-C) + \ln(1-C)^2} \right) .
\end{aligned}
\tag{S11}
$$

In Fig. S1 we see that numerical simulation of the model suggest that only the first two of these are evolutionary endpoints (respectively a zero fusion state and a diverging trajectory tending to large fusion rates).

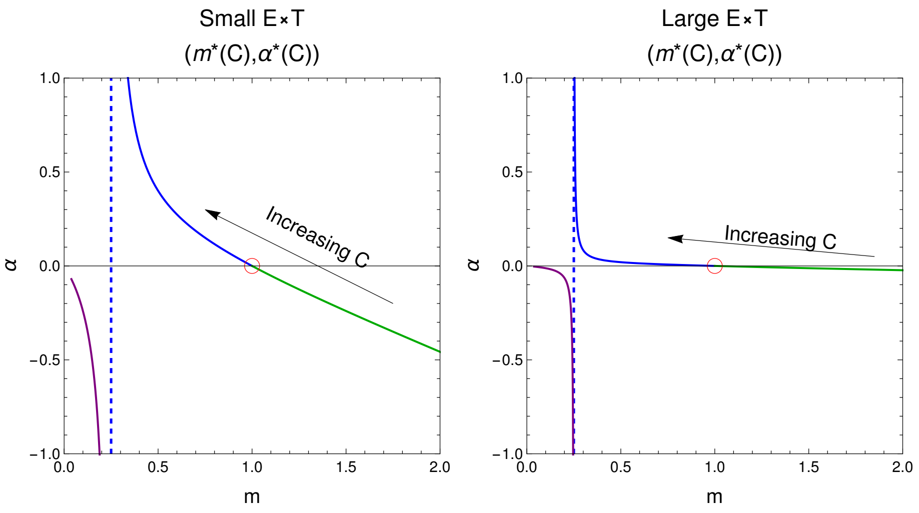

**Fig S2. Parametric plots of interior fixed point Eq. (S11) as a function of fusion cost,** $C$. The red circles show the location of the first point identified in Eq. (S11) and the vertical blue dashed lines show the $m = \beta/4$ manifold identified in Eq. (S11). The remaining solid lines show the location of the third interior point in Eq. (S11) when $1 - \mathrm{e}^{-1/2} > C$ (green), when $1 - \mathrm{e}^{-2} > C \geq 1 - \mathrm{e}^{-1/2}$ (blue), and when $C > 1 - \mathrm{e}^{-2}$ (purple). In both plots $\beta = 1$. In the left plot $ET = 2.5$, while in the right plot $ET = 50$.

#### S2.1 Interior Saddle Point

We begin by focusing on the third point identified in Eq. (S11), since it is the only point that varies with $C$. In Fig. S2 we show that this point is only biologically relevant if $1 - \mathrm{e}^{-2} > C \geq 1 - \mathrm{e}^{-1/2}$ (i.e. if the point exists in the positive quadrant). We now seek to determine the stability of this fixed point over this range and verify that it is indeed not attracting. Evaluating the Jacobian of the evolutionary dynamics given in Eq. (S10) at the interior fixed point, we obtain a matrix of the form

$$J = \begin{pmatrix} a & b \\ c & 0 \end{pmatrix} \tag{S12}$$

with

$$\begin{aligned} a &= -\frac{4\log(1-C)\left(\log^2(1-C)(2\log(1-C)+3)-2\right)}{3\beta^2}\,, \\ b &= \frac{2ET\log(1-C)(\log(1-C)+2)^2}{3\beta^2}\,, \\ c &= \frac{ET\log^2(1-C)(\log(1-C)+2)^2\log\left(\frac{6}{\log(1-C)+2}-3\right)}{3\beta^2(2\log(1-C)+1)}\,. \end{aligned} \tag{S13}$$

We further note that

$$a\begin{cases}=0 & \text{if} \quad \mathrm{C}=0\\ <0 & \text{if} \quad \mathrm{C}\in(0,1]\end{cases},$$
$$b=\begin{cases}=0 & \text{if} \quad \mathrm{C}=0\,\text{or}\,\mathrm{C}=1-\mathrm{e}^{-2}\\ <0 & \text{if} \quad \mathrm{C}\in(0,1]\backslash\left\{1-\mathrm{e}^{-2}\right\}\end{cases}, \tag{S14}$$
$$c=\begin{cases}=0 & \text{if} \quad \mathrm{C}=0\,\text{or}\,\mathrm{C}=1-\mathrm{e}^{-2}\\ <0 & \text{if} \quad \mathrm{C}\in(0,1-\mathrm{e}^{-2})\\ \in\mathbb{C} & \text{if} \quad \mathrm{C}\in(1-\mathrm{e}^{-2},1]\end{cases},$$

where $\mathbb{C}$ is the set of complex numbers.

Focusing on the interval $1-\mathrm{e}^{-2}>C\geq 1-\mathrm{e}^{-1/2}$ we see that $a$, $b$ and $c$ in Eq. (S12) are all strictly negative. The eigenvalues of $J$ are given by

$$\lambda_{\mathrm{s}}=\frac{a-\sqrt{a^2+4bc}}{2} \qquad \lambda_{\mathrm{u}}=\frac{a+\sqrt{a^2+4bc}}{2} \tag{S15}$$

from which we can infer $\lambda_{\mathrm{s}}<0$ and $\lambda_{\mathrm{u}}>0$, and thus the interior point identified in Eq. (S11) is a saddle. We next calculate the eigenvector associated with the stable direction for the saddle. We denote this vector by

$$V_{\mathrm{s}}=\begin{pmatrix}1\\ G\end{pmatrix} \tag{S16}$$

where $G$ can be understood as governing the gradient of the stable eigenvector, and is given by

$$G=\frac{2c}{a-\sqrt{a^2+4bc}}\,. \tag{S17}$$

#### S2.2 Zero fusion fixed point

We begin by noting that the point $(m^*,\alpha^*)=(\beta,0)$ is not a fixed point as one might understand it in a continuous dynamical system; it emerges instead because of the biological constraint $\alpha\geq 0$, which places a boundary at $\alpha=0$. Mathematically we find

$$\left.\begin{pmatrix}H_m(m,\alpha;\beta,C)\\ H_\alpha(m,\alpha;\beta,C)\end{pmatrix}\right|_{(m,\alpha)\to(\beta,0)}=\begin{pmatrix}0\\ -\frac{ET\left[1-(1-C)e^{-1/2}\right]}{2\beta}\,.\end{pmatrix} \tag{S18}$$

Because the dynamics are discontinuous at the boundary, one cannot employ a standard linear stability analysis as these leading-order terms in an expansion about the fixed point (which are precisely those given in Eq. (S18)) do not disappear [8].

In order to make analytic progress, we make the simplifying assumption that we can decompose the dynamics in $m$ and $\alpha$. Considering the dynamics of $m$ along the $\alpha=0$ boundary, it is clear that $(m^*,\alpha^*)=(\beta,0)$ is always stable in the $m$-direction. Meanwhile in the $\alpha$ direction it is clear that the leading order term for the dynamics of $\alpha$ in Eq. (S18) must be negative to prevent increases in $\alpha$ away from the fixed point at the boundary;

$$\begin{aligned}-\frac{ET\left[1-(1-C)e^{-1/2}\right]}{2\beta}&<0\\ \implies C&>1-\frac{1}{\sqrt{e}}\,.\end{aligned} \tag{S19}$$

We note that this transition point coincides with that where the interior fixed point crosses the boundary point (see Fig. S2). Simulations show that this is indeed a good analytic approximation for the critical value at which $(m^*, \alpha^*) = (\beta, 0)$ becomes stable (see Figs. 3 and S1). This can be understood intuitively from an analytic perspective by noting that the leading order term $ET\left[1 - (1-C)e^{-1/2}\right]/(2\beta)$ dominates the dynamics of small fluctuations in $\alpha$ away from the boundary in a series expansion of the dynamics.

#### S2.3 Trajectories on $m = \beta/4$ manifold

In order to make analytic progress, we take the crude limit $\alpha \to \infty$ to see what happens to the evolutionary dynamics in Eq. (5). We find

$$\left.\begin{pmatrix} H_m(m, \alpha; \beta, C) \\ H_\alpha(m, \alpha; \beta, C) \end{pmatrix}\right|_{(m,\alpha)\to(m,\infty)} = \begin{pmatrix} \frac{(\beta-4m)}{4m^2} \\ 0\,. \end{pmatrix} \tag{S20}$$

This suggests that at large values of the fusion rate, trajectories are attracted towards the manifold $m^* = \beta/4$.

Although the leading order terms in Eq. (S20) disappear when $m^* = \beta/4$, working in the limit $\alpha \to 0$ brings additional complications since the Jacobian is ill-defined in this limit. Essentially we find a zero eigenvalue associated with the fact that selection on $\alpha$ tends to zero in the $\alpha \to \infty$ limit. In order to make analytic progress, we again make the simplifying assumption that we can decompose the dynamics in $m$ and $\alpha$. We can see from Eq. (S20) that the manifold is always attracting in the $m$-direction for large $\alpha$. We next consider the dynamics of $\alpha$ on the $m^* = \beta/4$ manifold. We find

$$H_\alpha(m, \alpha; \beta, C)|_{(m,\alpha)\to(\beta/4,\alpha)} = -\frac{\left[1-(1-C)e^2\right]\beta\log\left(1+\frac{4ET\alpha}{\beta}\right)}{8ET\alpha^2(1-C)e^2 + 2\alpha\beta} \tag{S21}$$

which is positive (and therefore selects for increased fusion rates) if

$$C < 1 - \frac{1}{e^2} \tag{S22}$$

and is negative otherwise. We note that this transition point coincides with that where the interior fixed point crosses the $m^* = \beta/4$ (see Fig. S2). Simulations show that this is indeed a good analytic approximation for the critical value at selection for increased $\alpha$ flips from being positive ($C < 1 - e^{-2}$) to negative ($C > 1 - e^{-2}$), as illustrated in Figs. 3 and S1. We note that therefore when $C > 1 - e^{-2}$, the fixed point $(m^*, \alpha^*) = (\beta, 0)$ becomes the only evolutionary endpoint for the dynamics.

#### S2.4 Basin of attraction for zero fusion fixed point

We have seen in the previous sections that $(m^*, \alpha^*) \to (\beta/4, \infty)$ is the only possible evolutionary endpoint when $C < 1 - e^{-1/2}$ and that $(m^*, \alpha^*) = (\beta, 0)$ is the only possible evolutionary endpoint when $C > 1 - e^{-2}$. In this section we seek to characterise to which of these endpoints the population is attracted when $1 - e^{-2} > C > 1 - e^{-1/2}$, starting from zero fusion states $(m(0), \alpha(0)) = (m_0, 0)$.

A crude initial step would be to consider for what initial values of the daughter cell mass selection on $\alpha$ is positive, and therefore might lead to trajectories attracted towards increasing $\alpha$ along the $m^* = \beta/4$ manifold. We find

$$\begin{aligned} &H_\alpha(m, \alpha; \beta, C)|_{(m,\alpha)\to(m_0,0)} > 0 \\ &\implies m_0 < -\frac{\beta}{2\log(1-C)}\,. \end{aligned} \tag{S23}$$

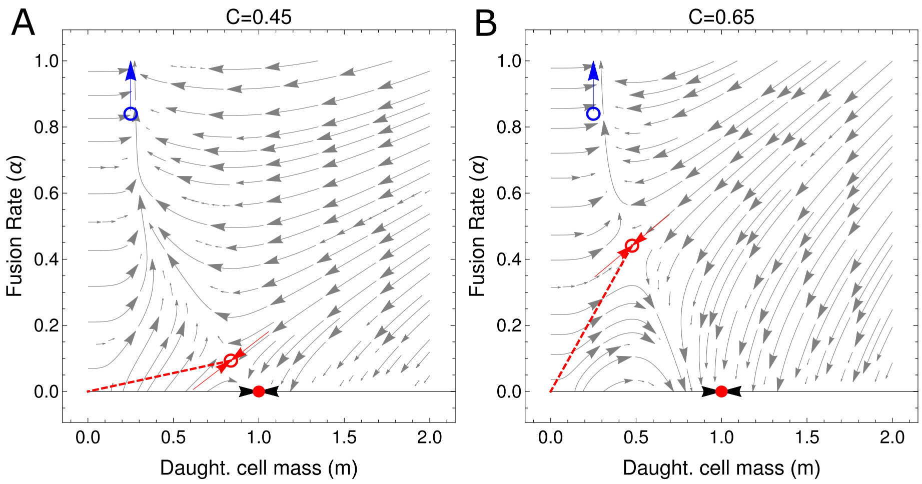

**Fig S3. Illustration of the calculation of the basin of attraction for the zero-fusion fixed point.** Blue arrow indicates the direction of the attracting $(m^*, \alpha^*) \to (\beta/4, \infty)$ manifold and red disk indicates the stable zero-fusion fixed point at $(m^*, \alpha^*) = (\beta, 0)$ (see Eq. (S11)). To which of these attractors the population evolves depends on whether the initial conditions on the $\alpha = 0$ boundary, $(m_0, 0)$, lie within the basin of attraction of $(m^*, \alpha^*) = (\beta, 0)$. We can approximate the separatrix for this basin of attraction using the stable eigenvectors of the interior saddle point (red arrows entering open circle, see Eqs. S11 and (S16)). In panel (A) with $C = 0.45$, the separatrix intersects the $\alpha = 0$ boundary; here high values of $m_0$ evolve towards $(m^*, \alpha^*) = (\beta, 0)$ and low values of $m_0$ evolve towards $(\beta/4, \infty)$. As $C$ is increased, the gradient of the stable eigenvectors eventually intersects the line connecting the origin and saddle fixed point (red dashed line) at $C = C_{\mathrm{Base}}$ (see Eq. (S24)). For $C > C_{\mathrm{Base}}$, we see in panel (D) that the entire $\alpha = 0$ boundary lies within the basin of attraction of $(m^*, \alpha^*) = (\beta, 0)$. Parameters are $\beta = 1$ and $ET = 2.5$.

We will see that this condition accurately predicts initial $m_0$ conditions that lead to selection for increased $\alpha$ along the $m^* = \beta/4$ manifold when $ET$ is large (such that the interior saddle fixed point is close to the $\alpha = 0$ boundary, see Fig. S2). However when $ET$ is small, this approach is misleading; as we see in Figs. 3 and S1, panels C and D, there are evolutionary trajectories that initially show an increase in $\alpha$ before eventually being attracted to the $(m^*, \alpha^*) = (\beta, 0)$. In order to fully characterize the values of $m_0$ that result in selection for large $\alpha$ we need to find the boundary for the basin of attraction of the $(m^*, \alpha^*) = (\beta, 0)$ fixed point, and in particular where this boundary intersects the $\alpha = 0$ axis. To do this we will develop a linear approximation for the separatrix defined using the stable eigenvectors of the interior saddle point.

Fig. S3 illustrates the conceptual details of the calculation. The stable eigenvector associated with the interior fixed point is given by Eq. (S16). We use a line with the same gradient $G$ passing through this interior fixed point as a linear approximation for the separatrix. We can then calculate the value $m = m_{\mathrm{Base}}$ at which this line crosses the $\alpha = 0$ boundary. Initial $m_0 > m_{\mathrm{Base}}$ then lie within the basin of attraction of $(m^*, \alpha^*) = (\beta, 0)$, while small $m_0 < m_{\mathrm{Base}}$ lie within the basin of attraction of $(m^*, \alpha^*) = (\beta/4, \infty)$.

We are particularly interested in the critical value of $C = C_{\mathrm{Base}}$ for which the basin of attraction comes to encompass the entire $\alpha = 0$ line (graphically, when the red arrows lie coincident with the dashed red line in Fig. S3). This will occur when the stable eigenvector gradient is equal to the gradient of the line intersecting the interior fixed

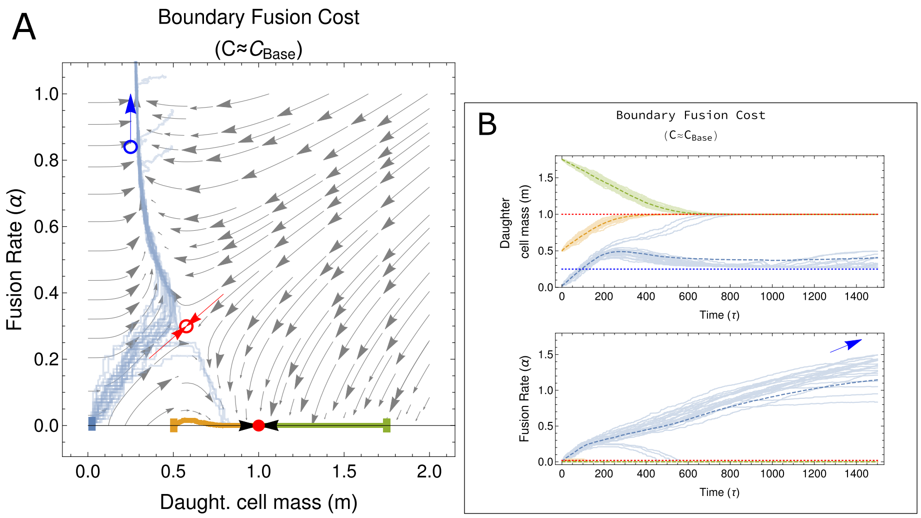

**Fig S4. Illustration of the dynamics near the $C_{\text{Base}}$ boundary.** Panel (A) shows the dynamics predicted by Eq. (5) in gray and the outcome of stochastic simulations for various initial conditions as colored lines; $m(0) = 0.02$ (blue), $m(0) = 0.5$ (orange), $m(0) = 1.75$ (green). Parameters are chosen such that the cost to cell fusion is only slightly greater than the critical cost $C_{\text{Base}}$ predicted by numerically solving Eq. (S24), with $C = 0.58$ and $C_{\text{Base}} = 0.573$. We can see that although the deterministic trajectories predicted by Eq. (5) do indeed lie within the basin of attraction for $(m^*, \alpha^*) = (\beta, 0)$, stochasticity in the numerical simulations can allow some trajectories to escape this basin of attraction for fusion costs close to the critical cost $C_{\text{Base}}$. Remaining parameters are $\beta = 1$ and $ET = 2.5$.

point and the origin, i.e. when

$$
\begin{aligned}
G &= \frac{\frac{\beta}{2\ln(1-C)}}{\frac{\beta}{ET}\frac{1+2\ln(1-C)}{2\ln(1-C)+\ln(1-C)^2}} \\
&= -\frac{2(2\log(1-C)+1)}{ET(\log(1-C)+2)} \\
&= \frac{2c}{a-\sqrt{a^2+4bc}},
\end{aligned}
\tag{S24}
$$

with $a$, $b$ and $c$ given in Eq. (S13). Note that in the limit of large $ET$, this condition simplifies to $G \approx 0$, which in turn implies $C = C_{\text{Base}} = 1 - e^{-2}$, while in the limit of small $ET$ we find $C = C_{\text{Base}} = 1 - e^{-1/2}$, as illustrated in Fig. 4 of the main text. Rearranging Eq. (S24) and eliminating roots outside the range $1 - e^{-2} > C > 1 - e^{-1/2}$, we obtain Eq. (13) in the main text.

In Fig. S4 we illustrate the evolutionary dynamics for costs to cell fusion near the critical value of $C$. The value of $C_{\text{Base}}$ obtained numerically is a good approximation of parameter regime at which the entire $\alpha = 0$ line lies within the basin of attraction of the fixed point $(m^*, \alpha^*) = (\beta, 0)$. However our simulations show that stochastically arising the order in which mutations are introduced and the finite mutational step sizes, $\delta m$ and $\delta\alpha$, can also play a role. Should these stochastic trajectories take the population across the separatrix, the population can escapte the zero-fusion basin of attraction. Thus near the critical point $C = C_{\text{Base}}$, only a fraction of the trajectories will evolve towards $(m^*, \alpha^*) = (\beta, 0)$, with the remainder evolving to high fusion rates.

#### S2.5 Implementation of Simulations

Here we detail the process of numerical simulation (see *Code availability statement*). We employ a multigenotype model whereby a mutation occurs at rate $\mu$. This rate is the inverse of the expected number of fusion processes (i.e. growth cycles) until the next mutation event. Before each successive mutation event, the number of fusion processes until the next mutation event is determined by generating a number from a $Geo(\mu)$ distribution and taking the inverse of that number. In an $S$ genotype model, if a given genotype $i$ has a trait value of $(m_i, \alpha_i)$ and has frequency $f_i$, then the mean population trait value is given by

$$\langle m \rangle = \sum_{i=1}^{S} m_i f_i \quad , \quad \langle \alpha \rangle = \sum_{i=1}^{S} \alpha_i f_i \tag{S25}$$

The simulations in Fig. 3 are repeated for 5500 mutation events. Below, we detail how we simulate the dynamics on each timescale.

### S3 Analysis of Evolutionary ODEs: Switching Environments

#### S3.1 Without Phenotypic Plasticity

We begin by considering a case in which the population switches between two environments, $\beta_1$ and $\beta_2$, but in which the population cannot evolve any phenotypic plasticity. In this scenario the population traits $m$ and $\alpha$ are fixed irrespective of the environment, such that the population can still be described by two traits $m$ and $\alpha$.

Switching between environments is modeled as a discrete stochastic telegraph process with a probability $\lambda_{i\to j}$ of transitioning from environment $i$ to $j$ at the beginning of each iteration of the life cycle (see Fig. 1). The time spent in each environment is distributed geometrically, with an average period $\tau_1 \approx 1/\lambda_{1\to 2}$ in environment 1 and $\tau_2 \approx 1/\lambda_{2\to 1}$ in environment 2. The probability of finding the population in either environment at long times is then given by $P_1 = \tau_1/(\tau_1+\tau_2)$ and $P_2 = \tau_2/(\tau_1+\tau_2) = (1-P_1)$.

In [4] we have shown that the evolutionary dynamics of a population in the scenario described above can be approximated by

$$\begin{aligned}\frac{\mathrm{d}m}{\mathrm{d}\tau} &= P_1 H_m(m,\alpha;\beta_1,C) + P_2 H_m(m,\alpha;\beta_2,C)\\ \frac{\mathrm{d}\alpha}{\mathrm{d}\tau} &= P_1 H_\alpha(m,\alpha;\beta_1,C) + P_2 H_\alpha(m,\alpha;\beta_2,C) \quad \text{with} \quad \alpha \geq 0\,.\end{aligned} \tag{S26}$$

This approximation works well so long at the environmental switching occurs at a rate equivalent to or faster than the timescale of mutation arrival [4].

In the present instance we are particularly interested in the case where $\alpha = 0$, as this gives us insight into possible initial conditions for $m_0$ in the scenario *with* phenotypic switching (see Main Text, *Mathematical Analysis and Results: Switching Environments with Phenotypic Plasticity*). We find

$$\begin{aligned}\frac{\mathrm{d}m}{\mathrm{d}\tau} &= P_1 H_m(m,0;\beta_1,C) + P_2 H_m(m,0;\beta_2,C)\\ &= P_1\left(\frac{\beta_1 - m}{m^2}\right) + P_2\left(\frac{\beta_2 - m}{m^2}\right)\,,\end{aligned} \tag{S27}$$

features a fixed point at

$$\begin{aligned}m^*_{\mathrm{BH}} &= \frac{P_1\beta_1 + P_2\beta_2}{P_1+P_2}\\ &= P_1\beta_1 + (1-P_1)\beta_2\,.\end{aligned} \tag{S28}$$

which is stable in the $m$-direction. This corresponds to a bet-hedging strategy in mass in which the population evolves to the average zero-fusion mass in each environment weighted by the probability of being in that environment.

#### S3.2 With Phenotypic Plasticity

We now consider the case in which the population switches between two environments, $\beta_1$ and $\beta_2$, but in which the population *can* evolve a phenotypically plastic response to each environment. In this scenario the population is now described by four traits, $(m_1, \alpha_1, m_2, \alpha_2)$, the control daughter cell mass and fusion rate in each respective environment. Environmental switching is again modeled as a telegraph process, as described in Section S3.1. We further assume that the population is initially in a phenotypically undifferentiated state $(m_1(0), \alpha_1(0), m_2(0), \alpha_2(0)) = (m_0, 0, m_0, 0)$ in which cell fusion has not yet evolved.

With costless phenotypic switching in which the population can immediately detect the environmental state, the evolutionary dynamics of the population in each environment is independent of that in the other. Equations for the dynamics in each environment can therefore be derived independently in the same manner as Section S1. The only modification is that the evolutionary timescale in a given environment is now moderated by the time spent in that environment;

$$\begin{aligned}
&\frac{\mathrm{d}m_1}{\mathrm{d}\tau} = P_1 H_m(m_1, \alpha_1; \beta_1, C), \quad \frac{\mathrm{d}\alpha_1}{\mathrm{d}\tau} = P_1 H_\alpha(m_1, \alpha_1; \beta_1, C) \quad \text{with} \quad \alpha \geq 0\,, \\
&\frac{\mathrm{d}m_2}{\mathrm{d}\tau} = P_2 H_m(m_2, \alpha_2; \beta_2, C), \quad \frac{\mathrm{d}\alpha_2}{\mathrm{d}\tau} = P_2 H_\alpha(m_2, \alpha_2; \beta_2, C) \quad \text{with} \quad \alpha \geq 0\,, \\
&m_1(0) = m_2(0) = m_0\,, \qquad \alpha_1(0) = \alpha_2(0) = \alpha_0 = 0\,.
\end{aligned} \tag{S29}$$

As noted in the main text, while the dynamics is decoupled in each environment, the trajectories in each environment are coupled through their shared initial conditions.

We are interested in the case where cell fusion evolves in one environment but not the other. We have seen that since the conditions in Eq. (14) are independent of $\beta$, the possibility of these two alternate outcomes in any environmental pair $(\beta_i, \beta_j)$ is contingent on intermediate costs, $C_{\text{Base}} > C > 1 - e^{-1/2}$; for higher costs cell fusion cannot evolve in either environment (see Section S2.4), while for lower costs cell fusion is the only evolutionary outcome (see Section S2.2).

In order to simplify our analysis and obtain analytic insight that is not possible when solving equations of the type given in Eq. (13) for the basin of attraction numerically, we work in the limit of large $ET$ (e.g. high density cells given ample opportunity for fusion). In this limit, the condition on $C$ for evolving cell fusion as a facultative stress response simplifies to $1 - e^{-2} > C > 1 - e^{-1/2}$, and the condition on the initial mass of the daughter cell simplifies to Eq. (S23). We will work within this limit for the remainder of this section.

###### S3.2.1 Initial condition $m_0 = \beta_1$ (population initially adapted to environment 1)

From Eq. (14) we see that $(m_0, \alpha_0) = (\beta_1, 0)$ is a stable state in environment 1 if $1 - e^{-2} > C > 1 - e^{-1/2}$ when $ET$ is large. We assume the population resides initially at in this zero-fusion state before being introduced to a novel harsh environment $\beta_2 > \beta_1$. This gives $m_2(0) = m_0 = \beta_1$. This initial condition leads to selection for high levels of cell fusion in environment 2 if Eq. (S23) is fulfilled;

$$\begin{aligned}
& m_2(0) < -\frac{\beta_2}{2\log(1-C)} \\
\implies\ & \beta_1 < -\frac{\beta_2}{2\log(1-C)} \\
\implies\ & \beta_2 > 2\beta_1 \log\left(\frac{1}{1-C}\right)\,.
\end{aligned} \tag{S30}$$

This means that the second environment has to be sufficiently harsh to trigger selection for cell fusion. Taking this condition together with those that keep the zero-fusion state stable in environment 1, we can see in Fig. 6(a) that facultative cell fusion as a stress response is possible when costs to cell fusion are intermediate and the novel environment is harsher than the first.

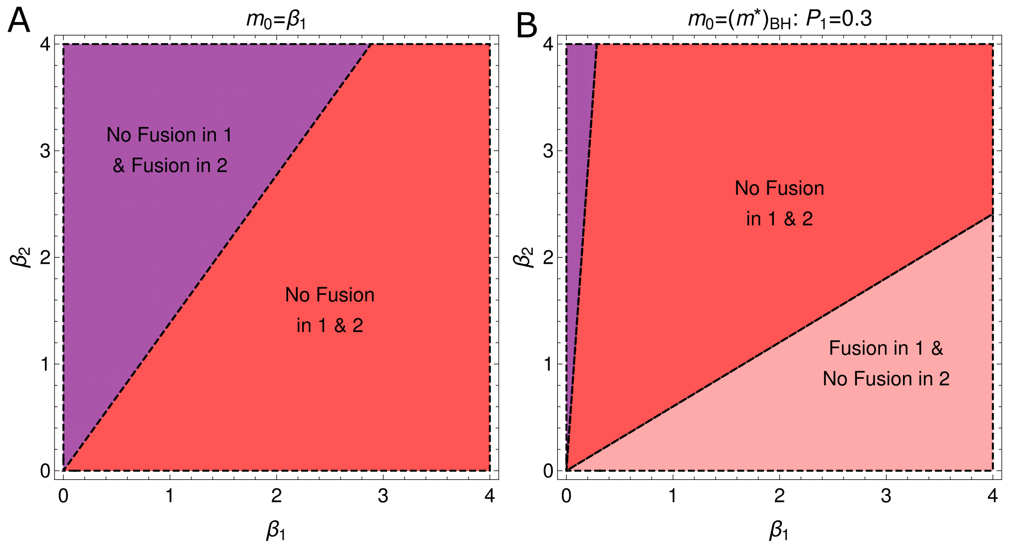

**Fig S5. Cell fusion can be observed in stressful environments.** Panels illustrate various potential outcomes of the evolutionary dynamics in the $\beta_1 - \beta_2$ plane in the large $ET$ limit for $C = 0.5$ (see also Fig. 6). Panel (A) shows outcomes for a population initially adapated to environment 1 (see Section S3.2.1) and Panel (B) shows outcomes for a population initially adapated to environment 1 (see Section S3.2.1). Purple regions show facultative cell fusion as a stress response to a harsh environment 2 ($\beta_2 > \beta_1$) and pink regions show facultative cell fusion as a stress response to a harsh environment 1 ($\beta_1 > \beta_2$). Red regions show parameters for which fusion does not evolve in either environment.

###### S3.2.2 Initial condition $m_0 = m^*_{\rm BH}$ (population initially adopting a bet-hedging strategy across environments 1 and 2)

We now consider the initial state $(m_0, \alpha_0) = (m^*_{\rm BH}, 0)$, with $m^*_{\rm BH}$ given in Eq. (S28) in which $\alpha$ has been fixed at zero. We now allow the daughter cell mass and the fusion rate to evolve as independent responses to each environment. In order to observe the evolution of binary cell fusion as a stress response we require trajectories starting at $(m_0, \alpha_0) = (m^*_{\rm BH}, 0)$ to evolve maintain zero fusion rates in one environment and towards large fusion rates in the other. We recall that when $ET$ is large, all initial conditions on the $\alpha = 0$ line eventually lead to selection for high fusion rates when $1 - e^{-1/2} > C$ and zero fusion rates when $C > 1 - e^{-2}$. We therefore focus on the interval between these two costs where cell fusion is a possible facultative outcome.

We begin by determining the conditions under which $(m_0, \alpha_0) = (m^*_{\rm BH}, 0)$ leads to zero fusion rates in environment 1. In the limit of large $ET$, we can use Eq. (S23) to find;

$$\begin{aligned} m^*_{\rm BH} &> -\frac{\beta_1}{2\log(1-C)} \\ \implies \beta_2 &> \beta_1 \left( \frac{P_1}{P_1 - 1} + \frac{1}{2(P_1-1)\log(1-C)} \right) ; \end{aligned} \tag{S31}$$

otherwise the population evolves to large fusion rates in environment 1. Similarly the condition under which the population evolves to zero fusion rates in environment 2 is given by

$$
\begin{aligned}
m^*_{\mathrm{BH}} &> -\frac{\beta_2}{2\log(1-C)} \\
\implies \beta_2 &< \beta_1 \left( \frac{2P_1 \log(1-C)}{2(P_1-1)\log(1-C)-1} \right),
\end{aligned}
\tag{S32}
$$

and otherwise the population evolves to large fusion rates in environment 2. In Fig. S5(B) we show some of the possible evolutionary outcomes. We again see that cell fusion in environment $j$ and not in environment $i$ is contingent on environment $j$ being comparatively harsher ($\beta_j > \beta_i$).

In Fig S6 we combine the results of Fig. S5(A-B) under the assumption $\beta_2 > \beta_1$ (i.e. environment 2 is harsher than environment 1). We then show in S7 that these analytic results are a good predictor of the results observed in simulations.

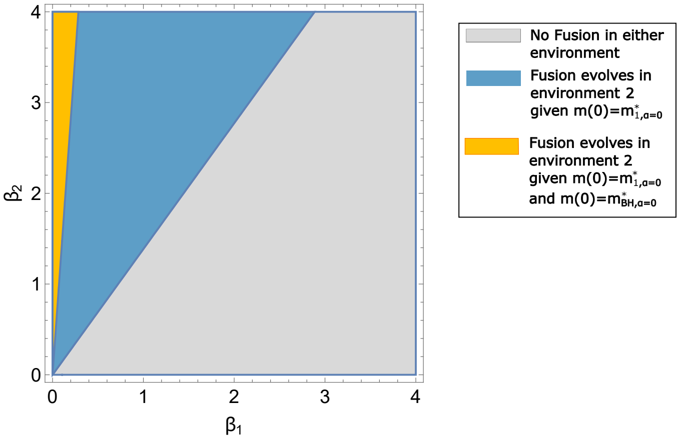

**Fig S6. Regions in the $\beta_1 - \beta_2$ plane where binary cell fusion evolves as a stress-response to environment 2**. This plot summarizes the outcomes in Fig. S5, again in the limit of large $ET$. For clarity we here further assume $\beta_2 > \beta_1$ so that fusion is restricted to evolving in response to environment 2. Fig. S7 illustrates a transect across the plane for $\beta_2 = 2$ and shows that numerical simulations support these analytic predictions.

#### S3.3 Implementation of simulations

We numerically simulate the evolutionary dynamics starting from the initial states as described in Figure 5 of the main text. To simulate the evolutionary dynamics for a population with initial conditions $(m_1(0), \alpha_1(0)) = (m_2(0), \alpha_2(0)) = (\beta_1, 0)$, we use a fixed value of $\beta_2$, and run the evolutionary dynamics as described in Section S2.5, for various values of $\beta_1$ (ranging from 0 to 4 in our simulations). To simulate the evolutionary dynamics for a population with initial condition $(m_1(0), \alpha_1(0)) = (m_2(0), \alpha_2(0)) = (P_1\beta_1 + (1-P_1)\beta_2, 0)$, we again run the evolutionary dynamics as described in the above sentence, using the same fixed value of $\beta_2$. The parameters used to run the evolutionary dynamics are $\delta = 1/50$, $\mu = 1/500$, $f_0 = 1/500$ and simulation run for $400/\mu$ growth cycles. All the other parameters are as in Figure S6 of main text. Running it for $2 \times 10^5$ growth cycles ($400/\mu$ growth cycles) provides sufficient time for the stochastic trajectory to reach the $m \approx \beta/4$ manifold in the alternate environment. At the 400th mutation event, we check whether the conditions $|m - \beta/4| < \delta$ and $\alpha > \delta$ are satisfied in order to determine whether the trajectory has reached the vicinity of the $m \approx \beta/4$ manifold. A numerical simulation is shown in Figure S7.

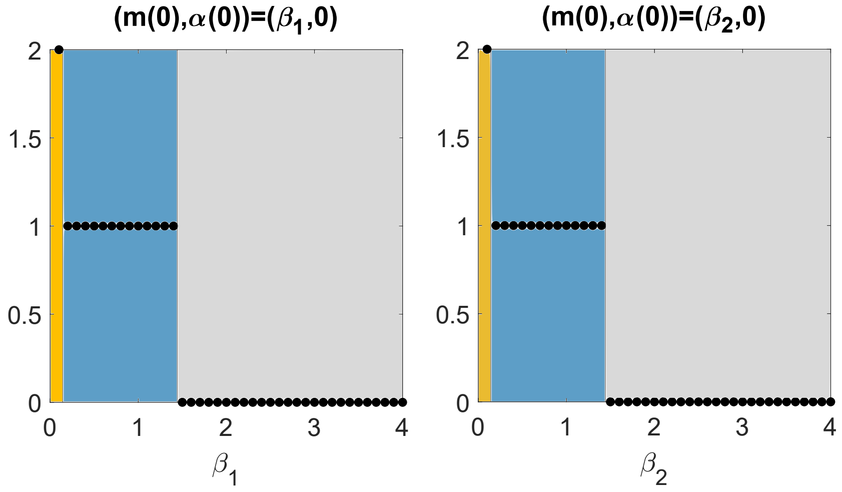

**Fig S7.** Black markers show numerical simulations of the evolutionary outcome in a switching environment with phenotypic plasticity. If the marker has a y-coordinate of 2, we evolve obligate fusion in environment $j$ given both $(m(0), \alpha(0)) = (\beta_i, 0)$ and $(m(0), \alpha(0)) = (P_1\beta_1 + (1 - P_1)\beta_2, 0)$ for $i \in \{1, 2\}$, $i \neq j$. If the marker has a y-coordinate of 1, we evolve obligate fusion in environment $j$ given $(m(0), \alpha(0)) = (\beta_i, 0)$ only (for $j \neq i$) and if the marker sits at 0, we evolve no fusion given either initial condition. Different coloured regions represent the behaviour predicted analytically predicted in Figure S6 of the main text. In left panel $\beta_2 = 2$ and $i = 1$ and in the right panel $\beta_1 = 2$ and $i = 2$. We see that the numerical simulations match the analytical predictions. The parameter conditions are $E = 100$, $T = 1$, $C = 0.5$, ($P_1 = 0.7$ if $\beta_1 > \beta_2$ and $P_1 = 0.3$ if $\beta_2 > \beta_1$), $\delta = 2 \times 10^{-2}$, $\mu = 2 \times 10^{-3}$ and $f_0 = 2 \times 10^{-3}$. Simulation is run for $2 \times 10^5$ growth cycles ($500/\mu$ growth cycles).

### S4 Parameters for simulations

In all the relevant figure panels, unless otherwise stated, the system parameters used are $E = 100$, $T = 1$ and $\beta = 1$ and the simulation parameters are $\delta = 10^{-2}$, $f_0 = 2 \times 10^{-3}$, $\mu = 10^{-3}$ (number of growth cycles)$^{-1}$. Each panel in Figure 3 of the main text has different fusion costs, with $C = 0.3$ in panel (A), $C = 0.45$ in panel (B), $C = 0.65$ in panel (C) and $C = 0.9$ in panel (D).

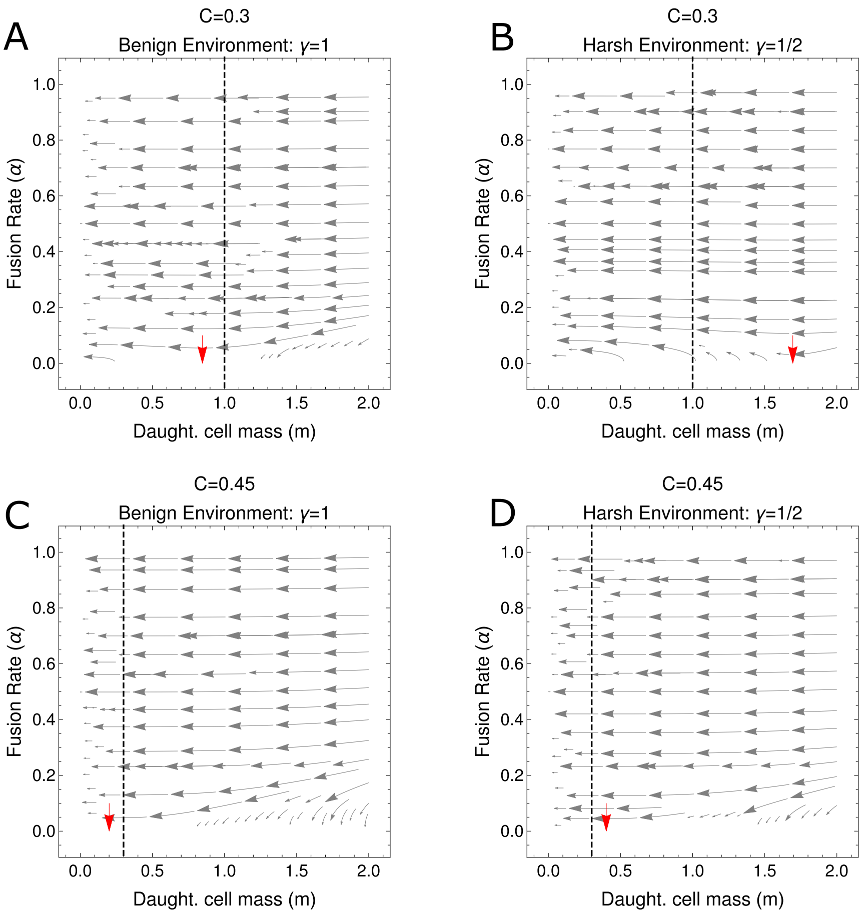

**Fig S8. Phase portraits for the evolutionary dynamics when using alternative survival function Eq. (S33).** Gray lines show the dynamics of Eq. (S34) and vertical dotted lines show the location of a boundary implemented at $m_{\mathrm{min}} = 1$ (panels (A) and (B)) and $m_{\mathrm{min}} = 0.3$ (panels (C) and (D)). In the limit $m_{\mathrm{min}} = 0$ all trajectories evolve towards the biologically unrealistic boundary $m = 0$. For $m_{\mathrm{min}} > 0$, trajectories evolve towards this new boundary. Red arrows indicate points on the $\alpha = 0$ boundary where selection on $\alpha$ switched direction (see Eq. (S35)); to the right of the arrow $(\mathrm{d}\alpha/\mathrm{d}\tau)|_{\alpha=0}$ is negative and to the left of the arrow $(\mathrm{d}\alpha/\mathrm{d}\tau)|_{\alpha=0}$ is positive. Note that for the $m_{\mathrm{min}}$ boundaries illustrated by black lines, selection on the boundary is negative in the benign environment and positive in the harsh environment (compare panel (A) with (B) and (C) with (D)). This illustrates the potential for stress induced binary cell fusion with the alternative survival function (see also Fig. S9). Remaining parameters are $ET = 10^3$.

### S5 Alternative Survival functions

We have primarily considered the dynamics of the model assuming a Vance survival function, $S(m; \beta) = \exp(-\beta/m)$. We now consider an alternative survival function

$$S(m; \gamma, m_{\mathrm{min}}) = \begin{cases} 1 - e^{-\gamma m} & \text{if} \quad m \geq m_{\mathrm{min}} \\ 0 & \text{if} \quad m < m_{\mathrm{min}} \,. \end{cases} \tag{S33}$$

The introduction of a minimum cell size required for survival, $m_{\min}$, is analogous to the decrease in cell survival below $m = \beta/4$ implicit in the Vance survival function (see Fig. 7 of the main text). We also note that $\gamma$ has an alternative interpretation relative to $\beta$; increased $\gamma$ indicates a more benign environment.

Following an analogous derivation as conducted in Section S1, we obtain

$$\begin{aligned}
\frac{\mathrm{d}m}{\mathrm{d}\tau} &= \frac{(1+m\gamma)(me^{m\gamma}+(1-C)E\alpha T)-e^{2m\gamma}(m+(1-C)E\alpha T)}{m\Big(e^{2m\gamma}(m+(1-C)E\alpha T)-(me^{m\gamma}+(1-C)E\alpha T)\Big)}\,, \quad m \geq m_{\min} \\
\frac{\mathrm{d}\alpha}{\mathrm{d}\tau} &= \frac{\big(1-C(e^{m\gamma}-1)\big)m\ln\Big(1+\frac{E\alpha T}{m}\Big)}{2\alpha\Big((1-C)E\alpha T+e^{m\gamma}(m+(1-C)\alpha TE)\Big)}\,, \quad \alpha \geq 0\,.
\end{aligned} \tag{S34}$$

Illustrative trajectories are shown in Fig. S8. We see that for $m_{\min} = 0$ there exists a selective pressure for the population to evolve to state producing infinitely many daughter cells of zero mass. This is clearly unrealistic biologically, and also leads to a mathematical breakdown of the evolutionary dynamical equations. We therefore focus on the case $m_{\min} > 0$.

We can see in Fig. S8 that the population will experience a strong selective pressure towards $m_{\min}$. Following an analogous approach to Section S3.2, we can investigate the dynamics of the population in a switching environment with costless phenotypic plasticity. In order to observe cell fusion as a facultative response, we require that at this point, selection for $\alpha$ is negative in one environment (i.e. there is an environment in which cell fusion held at zero) and positive in the other (i.e. there is an environment in which cell fusion selected for). The condition for positive selection on $\alpha$ in environment $i$ at $m = m_{\min}$ is

$$\begin{aligned}
&\left.\frac{\mathrm{d}\alpha}{\mathrm{d}\tau}\right|_{(m,\alpha)\to(m_{\min},0)} > 0 \\
&\qquad \implies m_{\min} < \frac{1}{\gamma_i}\ln\left(\frac{1-C}{C}\right)\,,
\end{aligned} \tag{S35}$$

as illustrated by the red arrows in Fig. S8. If this condition is fulfilled in one environment but not the other, then selection for fusion evolves as a facultative response. We can also see that in particular, cell fusion is selected only as a facultative stress response to detrimental environments that induce a reduction in cell survival rates (see Fig. S9). We also note that this condition cannot be fulfilled for any $m_{\min}$ if $C > 0.5$. In this high-cost scenario selection on $\alpha$ is strictly negative along the $\alpha = 0$ boundary. This places a more stringent upper bound on the costs to cell fusion for the survival function given in Eq. (S33) than are seen with the Vance survival function given in Eq. (14), which we recall permits the evolution of facultative cell fusion with costs of up to $C < 1 - e^{-2}$.

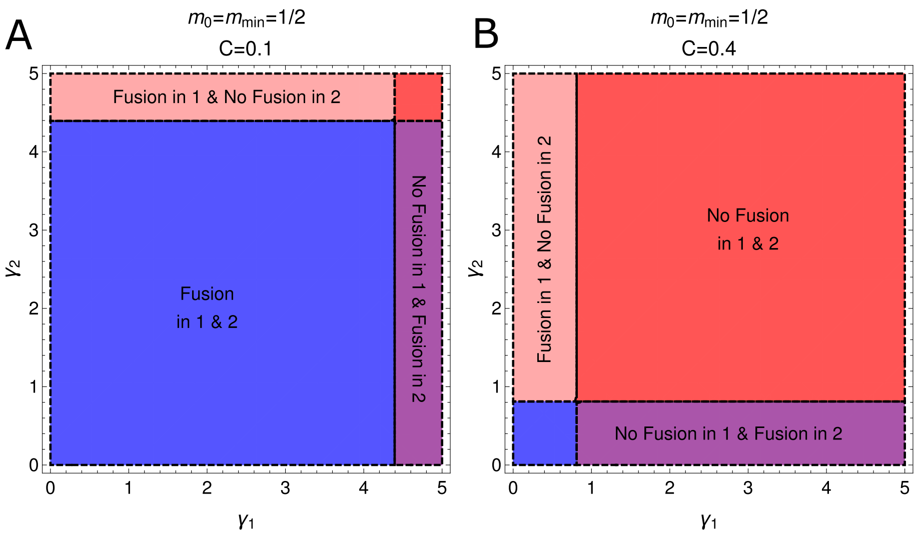

**Fig S9. Cell fusion can be observed in stressful environments when using alternative survival function Eq. (S33).** Recall that $\gamma_i > \gamma_j$ indicates that environment $j$ is harsher. Plots show the possible evolutionary outcomes of Eq.(S34) when $m_{\min} = 0.5$ and when $C = 0.1$ (panel (A)) and $C = 0.4$ (panel (B)). Smaller fusion costs increase the regions in which fusion evolves in both environments (blue) while larger fusion costs increase the regions in which fusion evolves in neither environment (red). Cell fusion can be observed as a facultative response to a harsh environment in the purple and pink regions. For costs exceeding $C = 0.5$ fusion cannot evolve in either environment.
